# Spatial Transcriptomics As Rasterized Image Tensors (STARIT) characterizes cell states with subcellular molecular heterogeneity

**DOI:** 10.64898/2025.12.18.695193

**Authors:** Dee Velazquez, Caleb Hallinan, Roujin An, Kalen Clifton, Jean Fan

## Abstract

Imaging-based spatially resolved transcriptomics (imSRT) technologies provide high-throughput molecular-resolution spatial characterization of genes within cells. Conventional analysis methods to identify cell-types and states in imSRT data rely on gene count matrices derived from tallying the number of mRNA molecules detected for each gene per segmented cell, thereby overlooking subcellular heterogeneity that can be useful in defining cell states. To take advantage of the molecular-resolution information in imSRT data and potentially identify cell-states based on subcellular heterogeneity, we developed STARIT (<u>S</u>patial <u>T</u>ranscriptomics <u>A</u>s <u>R</u>asterized <u>I</u>mage <u>T</u>ensors). STARIT converts transcripts within segmented cells in imSRT data into an image-based tensor representation that can be combined with deep learning computer vision models for downstream analysis. Using simulated and real imSRT data, we demonstrate that STARIT distinguishes transcriptionally distinct cell-types and further separates cell states based on subcellular transcript localization, which conventional gene count analysis fails to capture. By providing a standardized framework to encode subcellular molecular information in imSRT data, STARIT will enable deeper insights into subcellular heterogeneity and enhance the identification and characterization of cell-types and states that are overlooked by gene count representations.

**Author Summary:** Imaging-based spatial transcriptomics technologies can map the locations of individual RNA transcripts within cells, providing an opportunity to study how genes are spatially organized at subcellular resolution. However, most current analysis approaches simplify these data by counting transcripts per cell, reducing rich spatial information into cell-by-gene matrices and potentially overlooking biologically meaningful differences in subcellular transcript organization.

Here, we introduce STARIT (Spatial Transcriptomics As Rasterized Image Tensors), a Python framework that converts transcript coordinates into image-based tensor representations that preserve subcellular transcript organization. By combining these representations with computer vision feature extraction, STARIT enables analysis of subcellular transcript organization alongside conventional transcriptional abundance.

Using simulated and real imaging-based spatial transcriptomics datasets, we show that STARIT can recover expected cell-type-specific transcriptional heterogeneity while also potentially distinguishing cellular variation associated with transcript organization patterns that conventional count-based approaches fail to resolve. STARIT provides a general framework for incorporating subcellular transcript organization into spatial transcriptomics analysis and may facilitate the discovery of new cell states and biological variation overlooked by other analysis approaches.

## 1 Introduction

Recent advances in spatial transcriptomics technologies have enabled spatially resolved gene expression profiling for fixed cells in culture and thin tissue sections. In particular, imaging-based spatially resolved transcriptomics (imSRT) technologies, including MERFISH, seqFISH, Xenium, and others, enable researchers to pinpoint the location of individual mRNA transcripts for targeted genes within cells [1]. Unlike traditional fluorescence *in situ* hybridization, where a fluorophore’s color directly encodes the gene identity of a labeled molecule in an image, imSRT methods generally do not assign a detected transcript’s gene identity from a single image alone. Instead, imSRT methods decode a transcript’s gene identity based on multiple rounds of multiplexed imaging. The resulting output is typically represented as a table of decoded transcripts with associated spatial coordinates. To delineate cell boundaries, computational cell segmentation can be applied based on nuclear or membrane stains, total transcript density, or transcript composition [2,3].

Conventionally, analysis of such imSRT data has relied on assigning and tallying the number of transcripts detected for each gene per segmented cell (Figure 1A). This single-cell gene count matrix can then be used to identify transcriptionally distinct cell-types and states using dimensionality reduction and clustering approaches, analogous to conventional single-cell RNA-sequencing data analyses. However, these conventional analysis approaches inherently neglect the subcellular heterogeneity of these transcripts within cells such as potential nuclear, perinuclear, cytoplasmic, polarized, and other compartment-associated localizations. Such subcellular heterogeneity can play important roles in cellular function, as RNA localization regulates processes including local translation and intracellular signaling [4,5]. Likewise, disruptions in subcellular RNA localization have been linked to disease. For example, previous differential expression analysis of spatially-resolved RNA sequencing data has characterized the composition of synaptic RNAs in Alzheimer’s disease, implicating mislocalization of synaptic RNAs within neurons as a feature of a disease-associated cell-state [6]. While imSRT offers transcriptome-wide profiling to map subcellular RNA heterogeneity, computational frameworks capable of systematically incorporating such information remain limited. Considering subcellular heterogeneity in the analysis of imSRT data may enable the identification of functionally distinct cellular states previously unrecognized through conventional gene count analysis.

**Figure 1.**
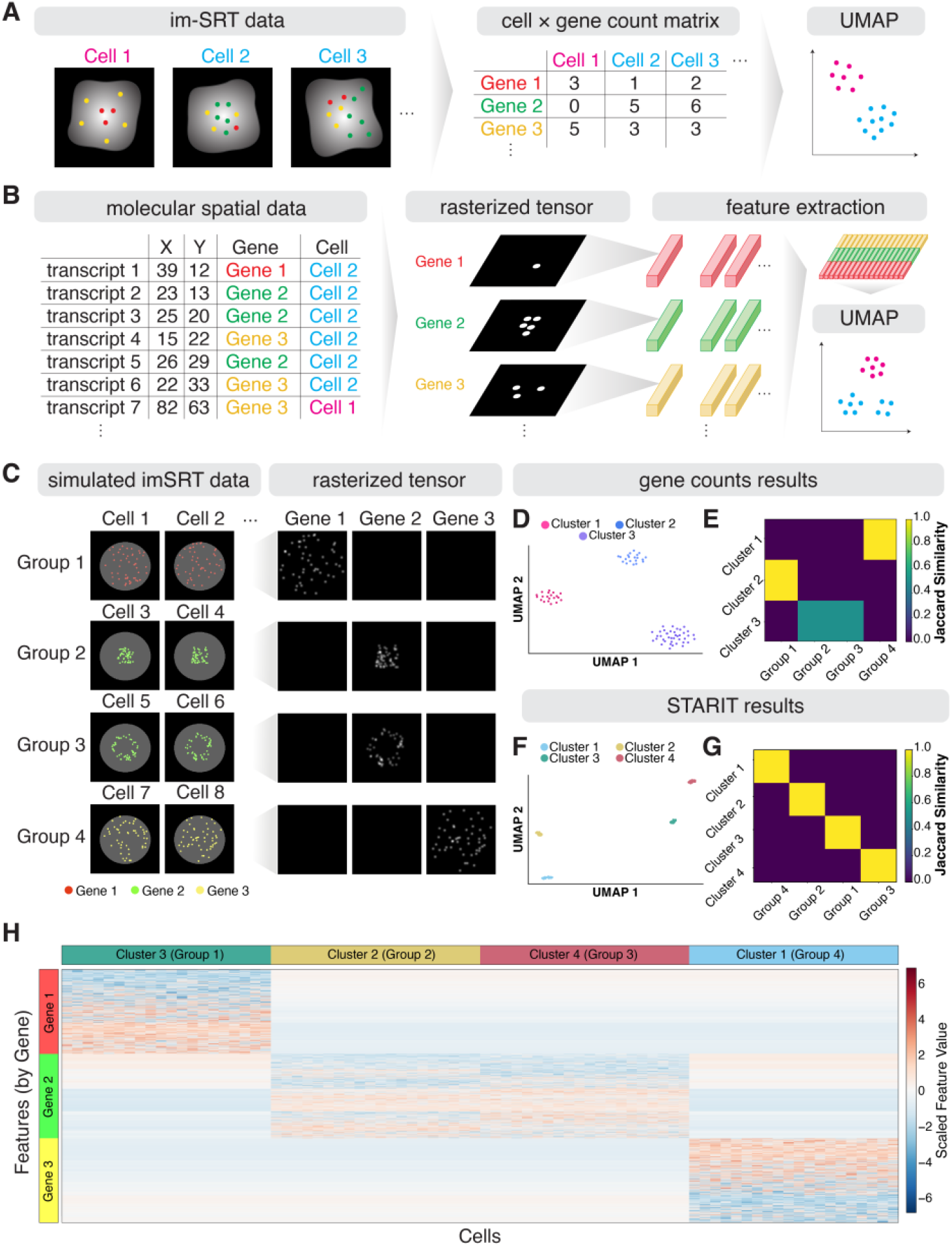
Overview of Spatial Transcriptomics As Rasterized Tensors (STARIT) (A) Schematic of conventional imaging-based spatially resolved transcriptomics (imSRT) data analysis pipeline. Detected transcripts are aggregated into a cell x gene count matrix, which is subsequently used for dimensionality reduction and clustering. Left: Three cells from imSRT data. Middle: Corresponding gene counts matrix. Right: Dimensionality reduction and clustering of imSRT identify distinct cell-types. (B) STARIT (Spatial Transcriptomics As Rasterized Image Tensors) workflow for imSRT data. Left: Molecular representation of imSRT data. Middle: Rasterized image representations for each gene in a cell. Right: Rasterized tensors are then input into a pretrained convolutional neural network (CNN; e.g., ResNet101) for spatial feature extraction. Extracted features are concatenated together to generate a total feature matrix that preserves both molecular and spatial information for downstream clustering and analysis. (C) Simulated imSRT data comprises four groups of cells (Groups 1–4) that exhibit distinct transcriptional and subcellular heterogeneity. Left: Example simulated cells from each group, showing expression of Gene 1 (red), Gene 2 (green), or Gene 3 (yellow). Groups are defined by distinct gene expression (Groups 1 and 4) or distinct subcellular localization patterns of the same gene (nuclear vs. perinuclear; Groups 2 and 3). Right: Rasterized tensor representations of each gene for one simulated cell per group. (D) UMAP visualization based on gene count analysis, with embedding colored by Louvain cluster assignment. (E) Jaccard similarity heatmap comparing ground-truth and gene count-based Louvain clusters. (F) UMAP visualization based on STARIT output, with embedding colored by Louvain cluster assignment. (G) Jaccard similarity heatmap between ground-truth and STARIT-based Louvain clusters. (H) Heatmap of scaled STARIT features grouped by gene channel, shown across the identified clusters. Rows denote extracted features, and columns denote cells.

As an alternative way to represent imSRT data to incorporate subcellular molecular spatial information and address this knowledge gap, we introduce STARIT (<u>S</u>patial <u>T</u>ranscriptomics <u>A</u>s <u>R</u>asterized Image <u>T</u>ensors) (Figure 1B). STARIT uses a previously developed rasterization strategy to preserve subcellular spatial molecular information by transforming dimensionless transcript coordinate-based measurements into pixel-based images thereby retaining organizational patterns [7]. Briefly, given the x-y coordinates of molecules within a segmented cell, STARIT creates one image per gene by encoding the presence of associated molecules within each image pixel as bits, which are then convolved with a Gaussian blur kernel to produce a smooth image that captures local molecular density. The pixel resolution is defined such that one pixel corresponds to one unit in the input coordinate system (Methods). Such use of rasterization to create image-based representations has previously been explored in spatial omics applications at the cellular level and hence STARIT represents an extension of these similar principles to the subcellular level [7,8]. All rasterized gene images for each cell are combined, resulting in a tensor representation. These tensor representations can then be fed into deep learning computer vision models, such as the pre-trained ResNet101 or UNI model, for feature extraction, resulting in a single cell feature matrix representation that captures the spatial distribution and concentration of molecules within cells. Given this single-cell feature matrix, conventional single-cell dimensionality reduction and clustering analysis pipelines can then be applied to identify clusters of cells with similar features, potentially reflective of distinct subcellular heterogeneity.

Using simulated and real imaging-based spatial transcriptomics datasets, we show that STARIT can distinguish cellular variation associated with transcript organization patterns that conventional count-based approaches fail to resolve while also recovering expected cell-type-specific transcriptional heterogeneity in real datasets.

## 2 Results

### 2.1 STARIT distinguishes cell states with subcellular molecular heterogeneity in simulated spatial transcriptomics data

To demonstrate the utility of STARIT for delineating cell states with subcellular molecular heterogeneity, beyond what conventional gene count clustering can capture, we first simulated imSRT data comprising of three transcriptionally distinct cell-types, each uniquely expressing one of three genes. For one cell-type, we further simulated cells such that they can be distinguished into two cell states based on nuclear versus perinuclear subcellular localization with no variation in expression magnitude between the two states. Specifically, we simulated cells of Group 1 to express Gene 1 uniquely, cells of Group 2 to express Gene 2 uniquely in a nuclear manner, cells of Group 3 to express Gene 2 uniquely in a perinuclear manner, and cells of Group 4 to express Gene 3 uniquely (Figure 1C). When we represent this simulated imSRT data as conventional gene counts and perform PCA dimensionality reduction and Louvain graph-based clustering, we are only able to identify three clusters of cells that correspond to the three simulated transcriptionally distinct cell-types (Figure 1D). This lack of correspondence between identified clusters and ground-truth groups can be further quantified via adjusted rand index (ARI) (score: 0.706) and the Jaccard similarity (mean: 0.833). While Cluster 2 corresponds perfectly to Group 1 cells and Cluster 1 to Group 4 cells, Cluster 3 contains a mixture of Group 2 and Group 3 cells. This overlap demonstrates that conventional representations of imSRT data and analyses are unable to distinguish between these two cell states, despite their distinct subcellular heterogeneity (Figure 1E). In contrast, when we apply STARIT, using the ResNet101 model for feature extraction followed by the same dimensionality reduction and clustering, we are able to identify four clusters of cells corresponding to the four simulated groups of cells representing cell-types and cell states with distinct subcellular heterogeneity (Figure 1F). This correspondence can be further quantified via ARI (score: 1.0) and in the on-diagonal Jaccard similarity (mean: 1.0) (Figure 1G). We further perform the Wilcoxon-rank sum test on the extracted features from ResNet101 and STARIT between cells in the four identified clusters as an analogous analysis to conventional differential expression testing. We identified significant features associated with the expected genes differentiating the four identified clusters, suggesting that the extracted features are capturing aspects of both cellular and subcellular molecular heterogeneity (Figure 1H).

Of note, while we used ResNet101, a feature extractor originally developed for natural images, for this analysis, STARIT is compatible with any downstream computer vision model for feature extraction. Although STARIT does not require the use of any specific feature extractor, feature extractor choice can influence which forms of spatial heterogeneity are emphasized. To demonstrate this, we generated an additional simulated dataset in which Groups 1 and 4 again uniquely expressed Gene 1 and Gene 3, respectively, while Groups 2 and 3 uniquely expressed Gene 2, now with left-versus right-polarized intracellular organization (Supplementary Figure 1A, Methods). Applying STARIT with ResNet101, we resolved four clusters corresponding to the four simulated groups, indicating that cell states defined by left-versus right-polarized transcript localization can be resolved (Supplementary Figure 1B). However, when we applied STARIT with UNI2-h, a feature extractor developed from a histopathology foundation model based on a vision transformer (ViT) encoder (ViT-H/14), to the same dataset, the analysis recovered three clusters, with reduced separation of the left- and right-polarized Groups 2 and 3 cells (Supplementary Figure 1C) [9]. Similar behavior was observed for CTransPath, another feature extractor for pathology images. In contrast, additional evaluated ResNet architectures including ResNet18, ResNet50, and ResNet152 more closely matched ResNet101 performance (Supplementary Table 1). This inability to distinguish left- and right-polarized groups may be expected because CTransPath, for example, is trained using frameworks that incorporate data augmentations such as random rotations and flips in learned visual representations [10]. Likewise, for UNI2-h, augmentation follows DINOv2 and other self-supervised ViT frameworks, which apply similar geometric transforms [9]. This result highlights that feature extractor choice may determine whether rotated and flipped spatial patterns such as left-versus right-polarization are preserved or diminished. Such sensitivity may be desirable when left-versus right-polarized spatial pattern is biologically meaningful, in the case of apical-basal organization, but undesirable when orientation reflects technical or arbitrary differences in cell positioning, for example. Thus, feature extractor selection should be guided by the biological question and expected sources of spatial variation in the dataset.

The simulated datasets thus far evaluate STARIT in settings where cell states are represented as discrete. However, cell states may vary continuously across developmental, environmental, or disease-associated trajectories [11,12]. We therefore sought to evaluate whether STARIT could capture continuous variation in subcellular organization. To model a continuous localization trajectory, we simulated cells with molecules sampled from either a circular nuclear region or a surrounding perinuclear annulus where the fraction of molecules assigned to the annulus increased monotonically from 0% to 100% across cells (Supplementary Figure 2A, Methods). As such, this simulation generated a gradual transition from nuclear to perinuclear transcript localization while holding total expression magnitude constant. STARIT preserved this gradual progression in various dimensional embedding spaces, suggesting that STARIT can capture continuous trajectories represented in a gradual transition of intracellular transcript organization (Supplementary Figure 2B).

Finally, a common challenge with current imSRT technologies, particularly in a tissue-setting, is that a whole cell may not always be captured in the associated thin tissue section. When a partial cell volume is captured, proportionally fewer transcripts may be detected. As such, we sought to evaluate whether STARIT remains robust when variable fractions of the cell volume are captured. We therefore generated additional simulated imSRT datasets to approximate partial cell volume capture in half the cells by systematically reducing molecule counts to represent 50%, 25%, and 12.5% volumetric capture variants, respectively (Supplementary Figure 3A-D, Methods). STARIT recovered the expected localization-defined groups in the 50% and 25% volumetric capture simulations, indicating robustness to moderate volumetric capture variation (Supplementary Figure 3E-F). However, performance degraded in the 12.5% volumetric capture setting, where sparse molecular sampling likely limited reliable recovery of subcellular localization patterns (Supplementary Figure 3G). These results suggest that STARIT can tolerate moderate reductions in transcript detection due to decreased volumetric capture. Based on these simulations, we recommend appropriate quality control filtering to remove cells based on low transcript abundance or segmented cell volume prior to applying STARIT.

In general, these simulation results demonstrate that STARIT enables an orthogonal approach for analyzing imSRT data to identify and characterize cell-states with diverse subcellular transcript localization patterns.

### 2.2 STARIT is sensitive to segmentation error, but can be remedied with gene-expression feature weighting

Having demonstrated the utility of STARIT on simulated data, we next sought to apply STARIT to real imSRT data. We applied STARIT to real imSRT data of the mouse cortex assayed by osmFISH, comprised of cell segmentation information and transcript spatial coordinates for 33 genes in 4,833 cells representing 31 cell-types (Supplementary Figure 4) [13]. Applying STARIT with ResNet101, we derived a 4,833 cell by 67,584 feature matrix and followed with PCA dimensionality reduction, Louvain graph-based clustering, and UMAP visualization (Supplementary Figure 5A). We identified 34 clusters using this approach and applied agglomerative clustering to obtain 31 clusters in an effort to recapitulate the original 31 cell-types identified by Codeluppi *et. al.* (Supplementary Figure 5B-C). However, we observed poor correspondence between our clusters to the original 31 cell-types (Supplementary Figure 5D). This lack of correspondence between identified clusters and ground-truth cell-types can be further quantified via ARI (score: 0.138) and in the on-diagonal Jaccard similarity (mean: 0.102). We hypothesized that this discrepancy could be driven by minor segmentation errors, noisy gene detection, or other technical artifacts, whereby misassignment of a few molecules would substantially impact our extracted image features.

To illustrate this effect, we used a simulated imSRT dataset comprising once again of four groups of cells, now augmented with a few misassigned non-cell-type-specific molecules representing minor segmentation errors and noisy gene detection (Methods, Figure 2A). When we apply STARIT to this simulated noisy imSRT data and perform PCA, Louvain graph-based clustering, and visualize in UMAP space, we can still identify four clusters of cells. However, these four clusters do not correspond to our four simulated groups of cells representing distinct cell-types and cell states but are rather impacted by the misassigned molecules (Figure 2B). This lack of correspondence can be further quantified via ARI (score: 0.582) and in the on-diagonal Jaccard similarity (mean: 0.616). As such, this confirms how the presence of misallocated mRNA molecules, such as from minor segmentation errors or noisy gene detection, can have an impact on STARIT.

**Figure 2.**
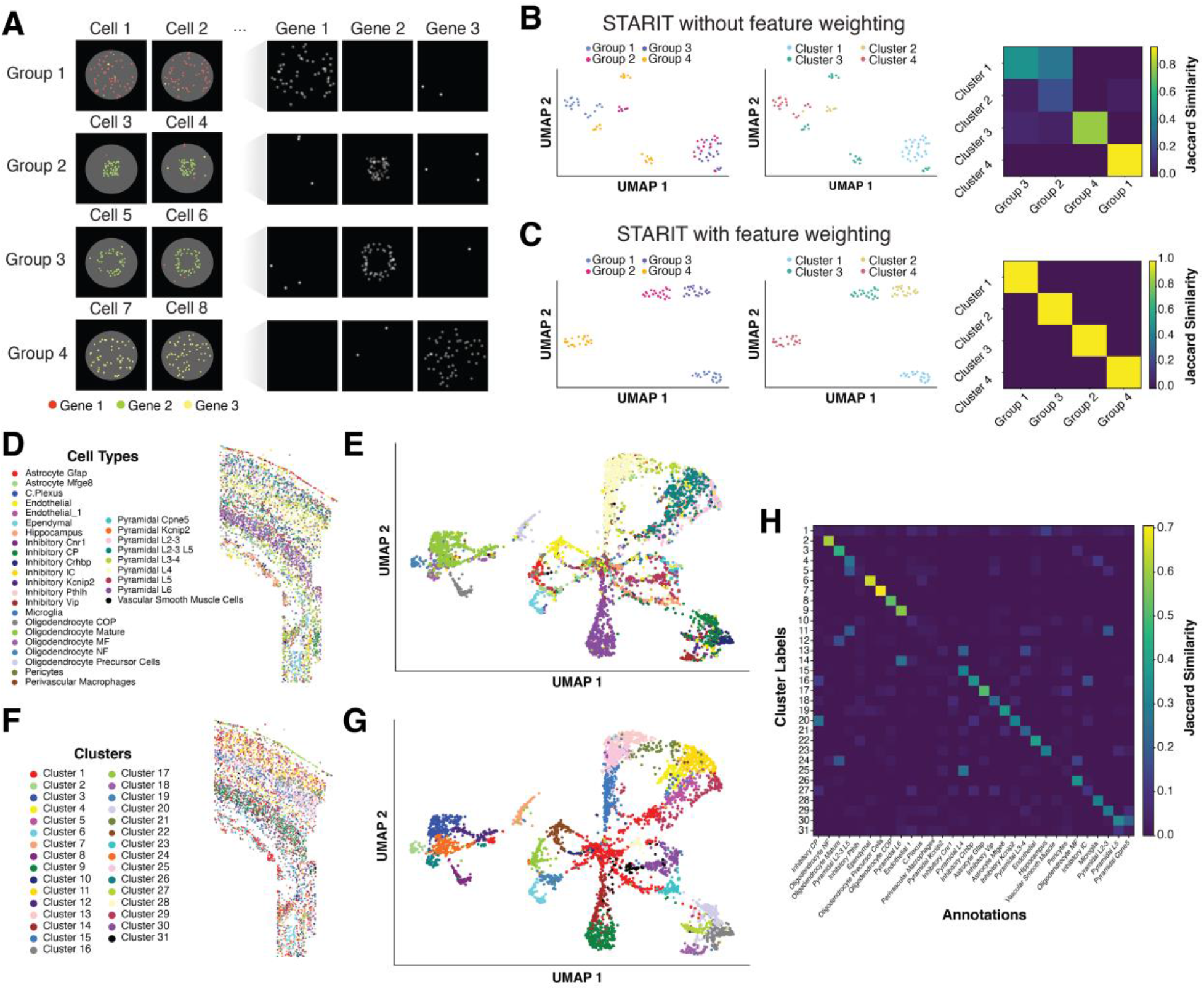
STARIT with feature weighting and application to osmFISH mouse cortex dataset. (A) Simulated imSRT data comprises four groups of cells (Groups 1–4) that exhibit distinct subcellular molecular heterogeneity. Left: Example simulated cells from each group. Groups are defined by distinct transcript expression and localization patterns across three genes (Gene 1, red; Gene 2, green; Gene 3, yellow). Groups 2 and 3 differ by nuclear versus perinuclear localization of Gene 2. All groups contain simulated noise across all genes. (B) STARIT analysis of simulated imSRT data with introduced noise, without feature weighting. Left: UMAP visualization of cells colored by ground-truth group annotation. Middle: UMAP visualization of cells colored by Louvain cluster assignment. Right: Jaccard similarity heatmap comparing ground-truth group annotations to Louvain clusters. (C) STARIT analysis of simulated imSRT data with introduced noise, with gene expression–based feature weighting. Left: UMAP visualization of cells colored by ground-truth group annotation. Middle: UMAP visualization of cells colored by Louvain cluster assignment. Right: Jaccard similarity heatmap comparing ground-truth group annotations to Louvain clusters. (D) Ground-truth cell-type annotations of osmFISH mouse somatosensory cortex dataset (Codeluppi *et al.*, 2018) in tissue space. (E) UMAP visualization of ResNet101-extracted STARIT features from the osmFISH mouse cortex dataset, colored by ground-truth cell-types. (F) STARIT-derived Louvain clustering of osmFISH mouse somatosensory cortex dataset in tissue space. (G) UMAP visualization of ResNet101-extracted STARIT features from the osmFISH mouse cortex dataset, colored by STARIT-derived Louvain clusters. (H) Jaccard similarity heatmap comparing ground-truth osmFISH mouse somatosensory cortex annotations to STARIT-derived Louvain clusters.

To address this issue, we introduced feature weighting to downweigh the contribution of such lowly detected genes in our STARIT analysis. Briefly, we multiply the extracted ResNet101 feature values for each gene in each cell by the min-max normalized expression magnitude of that gene in that cell. Such feature weighting effectively incorporates the prior expectation that more highly expressed genes are more important in defining a cell-type or state, downweighing lowly expressed genes that may have been detected due to minor segmentation errors and noisy gene detection. Using feature weighting with our simulated noisy imSRT data, we are able to identify four clusters of cells corresponding perfectly to the four simulated groups of cells representing distinct cell-types and cell states as quantified via ARI (score: 1.0) and on-diagonal Jaccard similarity (mean: 1.0) (Figure 2C). We further confirm that conventional gene counts clustering again identifies three clusters of cells, demonstrating that gene count clustering is more robust to such misassigned molecules but is still unable to resolve subcellular heterogeneity (Supplementary Figure 6). This establishes that while STARIT may be sensitive to minor segmentation errors and noisy gene detection, augmentation strategies such as weighting extracted features can be used to compensate to recapitulate expected results.

### 2.3 STARIT with feature weighting recapitulates expected cell-types

Having demonstrated that feature weighting can be an effective strategy to handle potential minor segmentation errors and/or noisy gene detection, we incorporate such feature weighting into our analysis of the osmFISH data. With feature weighting, we observe improved correspondence between our identified clusters and the original cell-type annotations, as reflected in the similar patterns in tissue and UMAP space as well as improved ARI with original cell types (score: 0.299) and higher on-diagonal Jaccard similarities (mean: 0.272) (Figs. 2D-H). This improved correspondence is consistent across anatomical regions (Supplementary Figure 7).

We do observe some new clusters unique to our STARIT analysis. For example, mature oligodendrocyte cells from the original osmFISH annotation are now split into clusters 3 and 24. As shown with our simulated data, such clusters may reflect distinct subcellular heterogeneity uniquely distinguishable through our STARIT analysis. We again used the Wilcoxon-rank sum test on the features between these two clusters to identify significantly differential features (Bonferroni-corrected p-value < 0.001) corresponding to the *Anln, Ctps, Gfap, Itpr2,* and *Plp1* genes’ STARIT representations. To evaluate whether our newly identified clusters are solely due to STARIT or could be attributable to inherent instability in gene count clustering analysis, we performed normalization, PCA dimensionality reduction, followed by Louvain graph-based and agglomerative clustering to identify 31 clusters from re-clustering on the original gene counts matrix. We were able to observe similar clustering results for the gene counts re-clustering as from our weighted STARIT analysis, reflected in the similar patterns in tissue and UMAP space (Supplementary Figure 8), with a comparable ARI score of 0.342 and on-diagonal Jaccard similarity of 0.295 (Supplementary Figure 8D). Again, these comparable correspondence is consistent across anatomical regions (Supplementary Figure 7).

These clustering results also included a splitting of mature oligodendrocyte cells (Supplementary Figure 8D). Consistent with this, when we used the Wilcoxon-rank sum test on the gene counts between the cells in the mature oligodendrocyte subclusters (cluster 3 and 24) from STARIT, we were able to identify the same genes (*Anln, Ctps, Gfap, Itpr2, Plp1*) as significantly differentially expressed (Bonferroni-corrected p-value < 0.001). These results suggest that STARIT identified new clusters that split annotated mature oligodendrocytes, potentially reflecting distinct cell states. However, re-clustering of gene counts could produce similar results, potentially indicating how subcellular localization changes can accompany subtle magnitude changes. Nonetheless, STARIT captures comparable aspects of transcriptional heterogeneity as conventional gene count analysis. Furthermore, analysis of STARIT-derived image features identified genes driving cluster differences comparable to differential expression with gene count clustering analysis.

### 2.4 STARIT distinguishes bacterial states with molecular heterogeneity in bacterial-MERFISH data

We next sought to apply STARIT to real imSRT data of cultured cells representing one cell-type to identify possible subcellular heterogeneity suggestive of distinct cellular states. We applied STARIT to imSRT data of *Escherichia coli (E. coli)* K-12 MG1655 bacteria, assayed by bacterial-MERFISH [14]. We focused on the 1000-fold volumetric expansion dataset, where the high expansion enabled reliable segmentation of bacteria confirmed by visual inspection. We performed quality control filtering to obtain 463 bacterial cells (Methods). As a proof-of-concept, we focused on one assayed operon, fliC-fliX. fliX is a post-transcriptional regulatory small RNA processed from the 3′ untranslated region (UTR) of the fliC transcript that base-pairs with fliC mRNA to regulate flagellar gene expression. The fliC gene encodes flagellin, a major flagellar filament subunit involved in bacterial motility [15]. Because fliC and fliX are co-transcribed as a single polycistronic mRNA molecule, they are detected as a single spatial unit within the bacterial-MERFISH probe design workflow [14,15]. We applied STARIT with ResNet101 to create a 463 cell by 2,048 feature matrix, followed by PCA dimensionality reduction and Louvain graph-based clustering. We identified 3 clusters of bacteria using this approach (Supplementary Figure 9A). Notably, these clusters of bacteria are not obviously distinguished based on fliC-fliX expression magnitude (Supplementary Figure 9B). Therefore, they would not have been readily found via conventional count-based analysis. When visualized in physical space, these 3 clusters appear distinct in their spatial rotation of flic-flix molecules (Supplementary Figure 9C). However, when visualizing the physical bacterial coordinates of these 3 clusters in culture, we observe that cells from different clusters exhibit distinct spatial rotations (Supplementary Figure 10). Therefore, in this example, the distinct spatial organization of fliC-fliX molecules is trivially driven by the rotation of the entire bacterium. Nonetheless, these results demonstrate that STARIT can resolve cells with distinct spatial heterogeneity including forms of rotations from well-segmented imSRT data of cultured cells.

To characterize potential biologically meaningful molecular spatial organizational heterogeneity that are not driven by the trivial rotation of the entire bacterium, we next sought to control for this rotational variation. To accomplish this, we standardized bacterial orientation across all 463 bacterial cells expressing fliC-fliX prior to rasterization (Methods). Briefly, for each bacterium, we centered all molecular coordinates and performed PCA in order to obtain the axis of maximal spatial variance, represented by the first principal component, to compute a rotation angle (θ) aligning the bacterium’s major axis to the x-axis. A corresponding counterclockwise 2-D affine transformation was then applied to all molecular coordinates to standardize orientation across cells (Supplementary Figure 11). To further minimize the impact of rotational variation, we applied STARIT with the UNI2-h feature extractor based on its previously demonstrated invariance to rotations via our simulations (Supplementary Figure 1C). We applied STARIT to these rotationally standardized molecular coordinates with UNI2-h to create a 463 cell by 1,536 feature matrix, followed by PCA and Louvain graph-based clustering. We identify 4 clusters of bacteria (Figure 3A). Again, these 4 clusters of bacteria are not obviously distinguished based on fliC-fliX magnitude (Figure 3B). When visualizing the physical bacterial coordinates of these 4 clusters in culture, we no longer observe that cells from different clusters exhibit distinct spatial rotations as expected (Supplementary Figure 12). We further perform the Wilcoxon-rank sum test on the UNI2-h features between cells in the 4 identified clusters to identify significant extracted features differentiating the 4 identified clusters (Supplementary Figure 13). However, because these extracted features are abstract, high-dimensional embeddings produced by the pretrained UNI2-h model rather than human-interpretable measurements, it remains unclear what exact aspect(s) of subcellular organization each feature captures, lacking direct biological interpretability.

**Figure 3.**
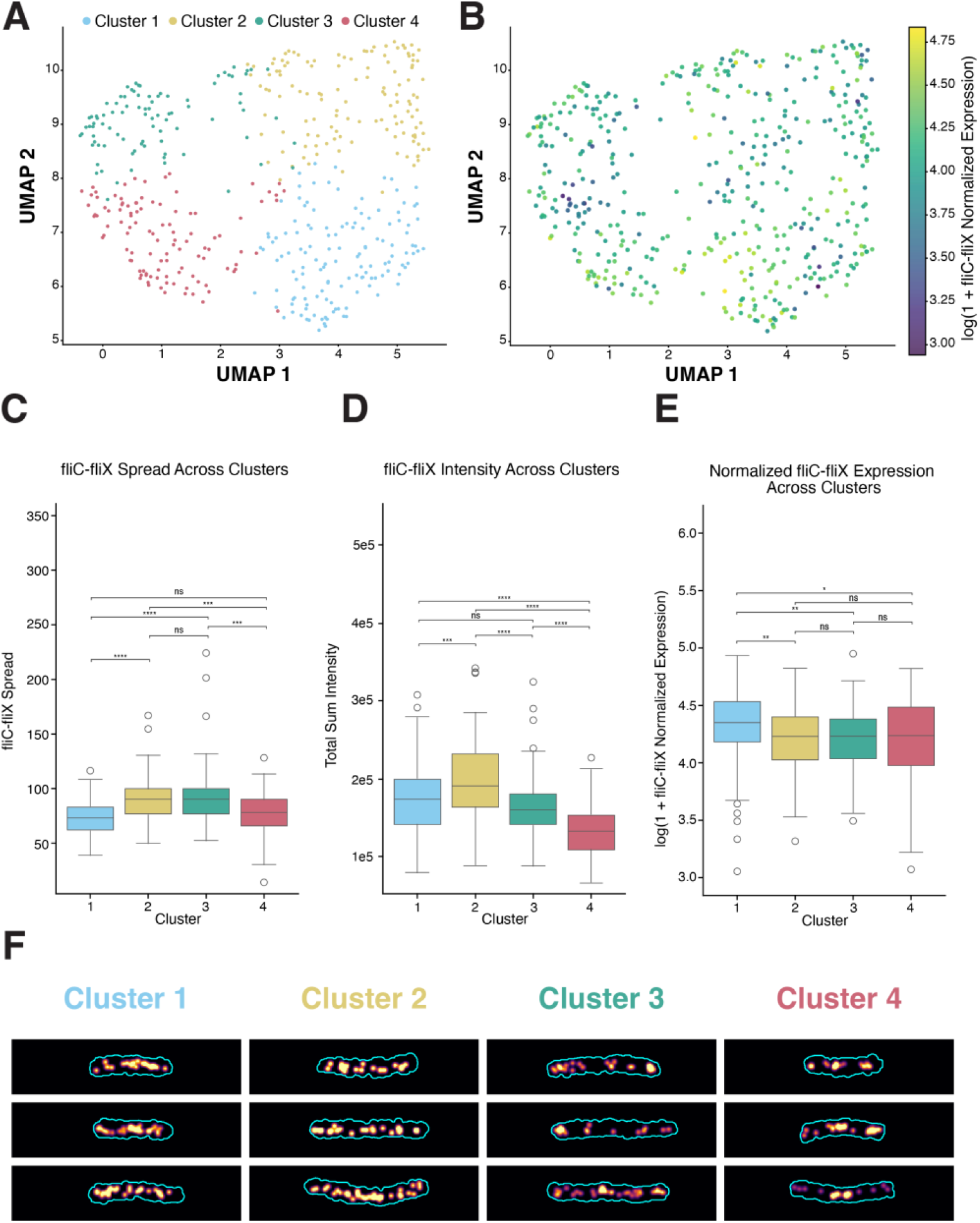
STARIT performance on standardized bacterial-MERFISH 1000X data. (A) Uniform Manifold Approximation and Projection (UMAP) of STARIT-derived UNI2-h features from rotationally standardized 463 *E. coli* cells based fliC-fliX operon localization, imaged with bacterial-MERFISH at 1000X volumetric expansion. (B) UMAP embedding of cells colored by log-transformed fliC-fliX gene expression levels. (C) Distribution of variation in normalized fliC-fliX gene expression across bacterial clusters. Expression levels normalized to total molecular counts per 1000 and log-transformed. Boxplots display median intensity (line), interquartile range (box), and outliers, with colors denoting cluster. Statistical significance determined by Mann-Whitney U test (Bonferroni corrected). (D) Distribution of spatial spread in X of fliC-fliX across bacterial clusters. Boxplots display median intensity (line), interquartile range (box), and outliers, with colors denoting cluster. Statistical significance determined by Mann-Whitney U test (Bonferroni corrected). (E) Distribution of total fliC-fliX signal across bacterial clusters. Boxplots display median intensity (line), interquartile range (box), and outliers, with colors denoting cluster. Statistical significance determined by Mann-Whitney U test (Bonferroni corrected). (F) Select examples of rasterized fliC-fliX transcript distributions for cells in clusters 1-4.

Therefore, to interpret the differences between the clusters identified by STARIT, we applied a supervised approach to extract a set of interpretable magnitude, morphological, and spatial organizational metrics from each bacterium (Methods, Supplementary Table 2). Specifically, we computed each bacterium’s normalized fliC-fliX expression magnitude and total operon expression from the original gene counts, the spatial fliC-fliX spread (in *x*) and cell length from the associated transcript coordinates, and the total fliC-fliX image intensity from STARIT where higher total intensity is associated with a more diffuse spatial distribution while lower total intensity is associated with a more punctated spatial organizational pattern (Methods, Supplementary Table 2). Pairwise two-sided Wilcoxon rank-sum tests on these supervised features revealed distinct combinations of magnitude, morphological, and spatial organizational properties among the 4 clusters (Figure 3C-E; Supplementary Figure 14). For example, Cluster 3 and Cluster 4 do not significantly differ in their normalized fliC-fliX expression magnitude (Figures 3E) and thus likely would not have been distinguished by conventional gene count clustering analysis. Cluster 3 does exhibit significantly greater fliC-fliX total intensity compared to Cluster 4 but also greater spread, cell length, and total operon count (Figures 3C-D; Supplementary Figure 14A-B). As such, the spatial heterogeneity in fliC-fliX distinguishing these two clusters may be trivially explained by cell size differences. In contrast, Cluster 2 and 3 neither significantly differ in their normalized fliC-fliX expression magnitude, spread, cell length, nor total operon count (Figure 3C, E; Supplementary Figure 14A-B). Yet, they differ significantly in fliC-fliX total intensity, with Cluster 2 cells exhibiting higher total intensities than Cluster 3 cells (Figure 3B). We can thus interpret Cluster 2 cells as having a more diffuse distribution of fliC-fliX transcripts throughout the bacterium and Cluster 3 cells as having a more punctated distribution of fliC-fliX transcripts (Figure 3F). Again, these difference in subcellular spatial organization would be missed by conventional gene count clustering and highlights the ability of STARIT to resolve cells with distinct subcellular spatial heterogeneity.

We note that these supervised metrics are highly correlated with some STARIT-derived UNI2-h extracted features (spread maximum Spearman = 0.61; intensity maximum Spearman = 0.64). This correlation suggests that the STARIT-derived UNI2-h extracted features, to some extent, capture these magnitude, morphological, and spatial organizational properties. Still, while fliC-fliX spread and total intensity are significantly different between our 4 identified clusters, these two supervised metrics do not as clearly delineate all cells across the 4 clusters as well as the UNI2-h extracted features (Supplementary Figure 15), suggesting that the bacterial subpopulations may be distinguished by a multivariate combination of features rather than by any two interpretable metrics.

Prior work has shown that bacterial mRNAs are not necessarily uniformly distributed within cells: mRNAs, ribosomes, and RNA polymerase can occupy distinct subcellular regions, and *E. coli* transcripts, including motility-related transcripts, can be asymmetrically enriched across cytosolic, membrane, and polar areas [16–18]. This polarization has been proposed to reflect localized translation of flagellar proteins, coordinating transcript localization with spatially constrained flagellar assembly [18]. In other bacteria, fliC translation is similarly controlled by intrinsic mRNA sequence or structure features that may be paired with secretion or assembly [19]. Given that fliC encodes the major structural component of the bacterial flagellar filament and fliX post-transcriptionally regulates flagellar gene expression, differences between dispersed and punctate fliC-fliX transcript distributions identified by STARIT may reflect proximity to flagellar assembly-associated machinery, differences in translational engagement, or other transcript organizational states. Because the bacterial-MERFISH data measures mRNA position but not ribosomes or flagellar proteins directly, there is no evidence of active localized translation directly. Nevertheless, the distinct fliC-fliX spatial localization patterns present in these clusters of bacterial cells despite overall similar normalized fliC-fliX expression magnitude suggests that STARIT can identify cell-states with biologically meaningful subcellular transcript localization patterns.

## 3 Discussion

Conventional representations of imSRT data, such as gene count matrices, inherently overlook spatial distributions of transcripts within cells, thereby inhibiting the identification of potential new cell states based on subcellular molecular heterogeneity. We developed STARIT to represent imSRT data as image-based tensors that preserve such subcellular molecular heterogeneity. On simulated imSRT data, STARIT combined with computer vision models for feature extraction followed by clustering analysis distinguished between cell states with distinct nuclear versus perinuclear molecular localization missed by conventional gene count clustering analysis. Likewise, in additional simulations, STARIT demonstrated robustness to partial cell volume capture and can further recover continuous trajectories. On real imSRT data, STARIT generally recapitulated expected cell-types in the mouse brain using osmFISH data and identified subcellular heterogeneity in *E. coli* using bacterial-MERFISH data that was not readily explained by expression magnitude alone.

Despite these advances, several limitations remain. As we have demonstrated, STARIT relies heavily on accurate cell segmentation and detection specificity, as such segmentation errors and noisy transcripts can bias image-derived features. To mitigate these effects, STARIT currently employs feature weighting to downweigh the contribution of image features derived from lowly expressed genes. However, such feature weighting may mask true biology driven by subcellular localization of lowly expressed genes. Alternative feature weighting based on other quality metrics beyond expression magnitude may be used in the future. In general, these results are consistent with recognized persistent challenges in cell segmentation and detection specificity for imSRT data analysis [20,21]. As imSRT technologies advance and segmentation quality improves for molecular resolution imSRT data, we expect the need for such feature weighting corrections to diminish.

Likewise, thin tissue sections currently used in imSRT introduce an additional source of variation because measurements may capture fully intact cells as well as partially sectioned cells with reduced volume. Our partial volume capture simulations suggest that STARIT can tolerate moderate transcript loss when the underlying localization pattern remains sufficiently represented. Still, sectioning through a cell may remove transcriptionally distinct structures to alter observed transcript localization patterns beyond transcript loss. Evaluating the robustness of STARIT under spatial truncation scenarios remains a direction for future work. Nevertheless, as with other single-cell and spatial analysis methods, these results demonstrate how applications of STARIT to thin tissue sections will require quality control filtering to remove cells based on low transcript counts or small captured cell volumes.

More generally, STARIT assumes that the observed two-dimensional transcript pattern snapshot contains enough information to approximate biologically meaningful subcellular organization. While this assumption may be weakened when only a small fraction of the cell volume is captured, ongoing advances in imSRT technology development toward applications in thicker tissue sections will strengthen its validity [22–25].

Moving beyond snapshots, many biological processes—such as disease progression, differentiation, and treatment response—progress continuously. Our nuclear-perinuclear continuum simulation suggests that STARIT-derived feature embeddings can preserve gradual transitions in subcellular transcript organization in addition to discrete states. Thus, evaluating STARIT on real imSRT datasets to discover such continuous transitions in subcellular organization along trajectories remains a potential direction for future exploration.

STARIT is designed as a general framework for coordinate-based imSRT technologies and can in principle be applied across technologies including but not limited to MERFISH, seqFISH, Xenium, CosMx, and related platforms. However, platform-specific technical properties may influence performance. For example, differences in segmentation strategy, transcript detection sensitivity, probe specificity, and off-target detection can alter the apparent subcellular distribution of transcripts [21,26,27]. As such, the robustness of STARIT to platform-specific technical variation represents an important future direction for determining whether image-based transcript localization features can consistently reveal biologically meaningful subcellular heterogeneity across platforms.

Feature interpretability also presents an ongoing challenge. As we’ve demonstrated, STARIT is compatible with any downstream computer vision model for feature extraction, but these models generally function as a “black box” with limited biological interpretability. For example, simply applying ResNet-based feature extraction to one rasterized image per gene for thousands of cells can generate a large feature matrix, with each feature potentially reflecting a mixture of signal related to transcript abundance, spatial spread, local density, shape, orientation, or potential technical artifacts. STARIT currently uses all extracted features for downstream dimensionality reduction and clustering without explicit feature selection before dimensionality reduction. While differential testing can identify features that significantly differ between identified clusters, as we have demonstrated, their biological meaning often remains open to interpretation. Likewise, while explainable AI approaches such as GradCAM and GAINext can provide coarse localization of model attention, they do not explain why specific features drive downstream cluster assignments [28,29]. In the bacterial-MERFISH analysis, supervised metrics helped characterize STARIT-derived clusters, but no single metric fully explained cluster identity. These results suggest that while supervised metrics can improve interpretability, they function as approximations of complex latent image features. We also anticipate that future incorporation of semi-supervised statistics regarding subcellular molecular organization could help further interpret cluster differences [30–32].

The choice of feature extractor is another important consideration. We primarily used ResNet101 as the feature extractor though we also evaluated alternative architectures, including smaller and larger ResNet models as well as domain-specific or foundation-model feature extractors such as CTransPath and UNI2-h (Supplementary Table 1). These comparisons suggest that feature extractor choice can influence downstream clustering and should be selected based on the biological question, data, and expected sources of variation. Computational cost is another practical consideration, particularly for large imSRT datasets. Because STARIT rasterizes and extracts features for every gene independently within each cell, runtime scales primarily with the number of cells and genes assayed. For example, applying STARIT to the full osmFISH dataset (4,833 cells, 33 genes) required approximately 2 hours for rasterization. By contrast, analyzing a targeted subset of the bacterial-MERFISH dataset (463 cells, 1 gene) completed rasterization in under 4 minutes. All reported runtimes reflect serial, non-parallelized execution on the hardware specifications described in Methods (4.16) and are intended to give readers a general sense of expected rasterization runtime. Exact runtimes will vary with computer specifications, parallelization strategy, if any, and dataset size. Downstream feature extraction runtime depends on the chosen computer vision model and its embedding dimensionality rather than on STARIT rasterization itself, and varied accordingly across our analyses: ResNet101 feature extraction required over 2 hours for the full osmFISH dataset and approximately 13 minutes for the bacterial-MERFISH subset, while UNI2-h feature extraction of the bacterial-MERFISH subset required just over 2 minutes. As the throughput of imSRT data improves, applying STARIT across all cells and genes in imSRT datasets with hundreds to thousands of targets may become computationally demanding. Parallelization and use of high-performance computing clusters may alleviate runtime concerns. Alternatively, we suggest a strategy in which conventional gene-count clustering is first used to identify cell types of interest, and STARIT is then selectively applied only to cells and/or gene subsets of interest, for example within a single annotated cell-type, to investigate potential subcellular heterogeneity. This approach can reduce computational cost while retaining STARIT’s ability to reveal subcellular heterogeneity within specific populations of interest.

A related limitation of ResNet architectures is rotational and scale sensitivity. Conventional convolutional neural networks, including ResNet models, are not inherently invariant to image orientation or scale. In the bacterial-MERFISH analysis, the original orientation of cells influenced extracted image features, highlighting both a limitation and a diagnostic property of image-based representations. To address this, we rotationally standardized cellular orientation before rasterization. Additional robustness could be achieved at the feature extraction step itself, for example by fine-tuning the existing pretrained feature extractor with random rotation and scale augmentations, utilizing contrastive learning to encourage augmented views of the same rasterized gene images within cells or cell states to produce similar feature vectors, or adopting rotation-invariant architectures or equivariant neural networks as feature extractors [33–35]. These strategies may be particularly important for tissues with polarized structures, such as epithelia with apical–basal organization, where rotation may represent either a confounder or a biologically meaningful axis depending on the analysis goal.

Although ResNet101 performed well in our analyses, it was originally trained on natural images and is not specifically optimized for sparse molecular images or subcellular transcript localization. As imSRT datasets continue to grow in scale and resolution, there is an opportunity to develop foundation models specifically for subcellular molecular imaging data. Existing histology and microscopy foundation models may provide useful general-purpose image features, but they were not trained specifically to represent sparse transcript-coordinate images or subcellular molecular localization patterns [9,10,36]. A foundation model trained directly on large-scale imSRT data across tissues, platforms, genes, and perturbations could learn representations better suited to molecular density, localization, polarity, and compartmental organization. Such models could improve feature robustness, reduce dependence on natural-image pretrained neural networks, and potentially support more interpretable representations of subcellular molecular organization in the future.

Although STARIT currently uses rasterized image representations, rasterization is only one strategy for encoding subcellular transcript organization. Alternative representations may directly model transcript coordinates without converting them into pixels through rasterization. Graph-based approaches have already been used in spatial omics to model spatial organization at the tissue or cellular-neighborhood level [37–39]. Likewise, such graph-based representations of transcript locations within cells has been demonstrated to improve cell type annotation [40] and cell segmentation [41]. Alternative point-cloud-based representations have also been developed to classify subcellular RNA localization patterns [42]. Future work comparing these representations can help determine which best captures biologically relevant information for specific analytical objectives.

Looking forward, by providing an alternative image-based representation of imSRT data, STARIT opens opportunities to integrate image-based representations of imSRT data with many downstream applications. Emerging frameworks such as scPortrait, a scverse-native toolkit that standardizes single-cell microscopy images in an AnnData-compatible format, illustrate how STARIT and related rasterization workflows could be wrapped into interoperable packages within broader computational infrastructure [43]. More generally, as imSRT technologies advance and cell segmentation improves, and likewise computer vision models advance and interpretable feature extraction improves, we anticipate that image-based representations of spatial transcriptomics will become a useful approach for resolving subcellular organization and for generating biologically meaningful interpretations. Overall, we anticipate that STARIT will enhance the identification and characterization of novel cell-types and states by capturing subcellular molecular heterogeneity, providing deeper insights into cellular organization and functional diversity that were overlooked by gene count analysis methods alone.

## 4 Materials and Methods

### 4.1 Spatial Transcriptomics As Rasterized Image Tensors (STARIT)

STARIT converts transcript-level spatial coordinates from imSRT data into rasterized density image representations that preserve subcellular molecular organization for each segmented cell. STARIT takes three primary inputs: (i) an axis-aligned bounding box (bounding_box) and the (ii) x and (iii) *y* coordinate vectors of all detected transcripts you wish to rasterize for a given cell. To maintain the original cell aspect ratio, a bounding box helper function outputs the calculated minimum and maximum spatial extrema of the molecular coordinates assigned to a cell (min*_x_*, max*_x_*, min*_y_*, max*_y_*). The box bounded by these four values is then expanded by a factor called expand (default 1.1) to provide a small margin for the rasterized image. The bounding box image grid is then defined at a desired pixel resolution size (default 1.0, in the same units as the input coordinates). From there, each gene molecule is then rendered to the canvas grid by convolving a normalized Gaussian kernel, blur (default 1.0), centered at (*x*,*y*). The pixelated image is created by summing contributions from all gene molecules in the cell. The result is STARIT producing one PNG image per gene per cell. When a gene has no molecules, a blank black image of the canonical size is saved to preserve the tensor shape.

### 4.2 Image Preprocessing and Feature Extraction Workflow

Prior to model input, each rasterized gene image was converted to RGB before the image was resized to 224×224 pixels with aspect-ratio preservation and padding. Each image was proportionally scaled to fit within a target bounding box of 224×224 pixels, determined by the limiting dimension:

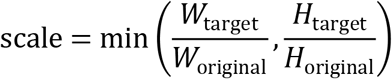

Images were downscaled using Lanczos interpolation. The resulting resized images were then symmetrically padded with black pixels (zero-padding) to fulfill the target dimensions, ensuring the original scaled images remained centered.

Images were normalized using ImageNet preprocessing parameters, with channel means of [0.485, 0.456, 0.406] and standard deviations of [0.229, 0.224, 0.225]. Unless otherwise specified, feature extraction was performed using pretrained computer vision models with the final classification layer removed.

For ResNet-based analyses, each gene-specific image was passed independently through the pretrained model. ResNet18 produced a 512-dimensional embedding per gene image, whereas ResNet50, ResNet101, and ResNet152 produced 2,048-dimensional embeddings per gene image. These per-gene embeddings were concatenated across all genes to generate one cell-level feature vector. For example, in simulated datasets with three genes, ResNet101 generated a 6,144-dimensional feature vector per cell. For the osmFISH dataset with 33 genes, ResNet101 generated a 67,584-dimensional feature vector per cell.

We also evaluated non-ResNet feature extractors, including CTransPath and UNI2-h. CTransPath produced 768 features per gene image whereas UNI2-h produced 1,536 features per gene image in the simulated analyses, with the resulting feature matrix dimensions reported in Supplementary Table 1. For all feature extractors, the resulting cell-by-feature matrix was used as input for dimensionality reduction, clustering, and visualization.

### 4.3 Gene expression-based feature weighting

To reduce the influence of image features derived from sparsely detected or potentially noisy genes, we implemented gene expression-based feature weighting. For each dataset, the corresponding cell-by-gene count matrix was min–max scaled per gene across cells. For each cell and gene, the image-derived feature vector for that gene was multiplied by the corresponding scaled gene count value. Weighted gene-specific feature vectors were then concatenated across genes to form a weighted cell-by-feature matrix.

This strategy preserves the image-derived spatial features for highly detected genes while downweighting features from genes with low or absent expression in a given cell. Feature weighting was applied to selected noisy simulated analyses and to the osmFISH analysis, as indicated in Supplementary Table 1.

### 4.4 Simulated imSRT Datasets

All simulated datasets were generated using Python scripts with fixed random seeds. Each simulated dataset was exported as a transcript-level coordinate table containing x-position, y-position, gene identity, cell identity, and group identity where applicable. A corresponding cell-by-gene count matrix was also generated for conventional gene-count analysis. Unless otherwise specified, each dataset consisted of 80 cells organized into four groups of 20 cells each, with three possible genes: gene 1, gene 2, and gene 3. Simulated cells were represented on a 224×224 coordinate canvas centered at (112, 112), with a circular cell radius of 75 pixels.

#### 4.4.1 Nuclear-perinuclear localization-marker simulation dataset

To evaluate whether STARIT could distinguish cell states defined by subcellular transcript localization under controlled conditions, we generated a fully synthetic simulated imSRT dataset known as simulated dataset #1 (Supplementary Table 3). The dataset simulated four groups of a known ground-truth spatial organization, each containing 20 circular “cells” represented within a 224×224 pixel canvas grid. For each cell, we simulated the spatial locations of transcripts for three genes, with each gene exhibiting a distinct, group-specific spatial pattern. Specifically, we generated 50 transcript coordinates for gene-group pairs defined to be expressed in that group. In Group 1 (gene 1) and Group 4 (gene 3), transcripts were uniformly sampled from anywhere within the full circular cell (radius = 75 pixels), resulting in a random cell-wide spatial pattern. In Group 2 (gene 2), transcripts were restricted to a smaller central region (radius = 25 pixels), producing a spatial pattern that reflects the nuclear space. In Group 3 (gene 2), transcripts were sampled from an annular region between circle radii of 25 and 45 pixels, yielding a perinuclear ring-like spatial pattern. For nuclear and perinuclear sampling, the nuclear region was defined as a disk centered at (112, 112), and the perinuclear compartment was defined as an annular region surrounding the nuclear region. Molecules were sampled uniformly within the allowed region for each group. Genes not active in a given group had zero molecules and were represented by blank black image channels after rasterization. The final simulated dataset consisted of transcript-level x, y coordinates annotated with gene, cell, and group identity, along with the derived traditional cell-by-gene count matrix for downstream evaluation and comparison.

#### 4.4.2 Left-right polarized localization-marker simulation dataset

To evaluate whether STARIT could distinguish additional forms of subcellular spatial organization beyond nuclear and perinuclear localization, we generated a left-right polarized localization-marker simulation, known as simulated dataset #2 (Supplementary Table 3). Groups 1 and 4 again uniquely expressed gene 1 and gene 3, respectively, with molecules sampled uniformly from the full circular cell. Groups 2 and 3 both expressed gene 2, but with distinct polarized localization patterns. In Group 2, gene 2 molecules were restricted to one side of the cell, whereas in Group 3, gene 2 molecules were restricted to the opposite side. This simulation was used to evaluate whether feature extractor choice influenced the recovery of orientation-dependent intracellular transcript patterns. The four simulated groups each contained 20 circular “cells” with 50 expressed transcripts of their respective gene represented within a 224×224 pixel canvas grid. Genes not active in a given group had zero molecules and were represented by blank black image channels after rasterization. The final simulated dataset consisted of transcript-level x, y coordinates annotated with gene, cell, and group identity, along with the derived traditional cell-by-gene count matrix for downstream evaluation and comparison.

#### 4.4.3 Simulated nuclear-perinuclear localization-marker dataset with sparse noise

To evaluate the robustness of STARIT and STARIT sensitivity to low-level background signal, we generated a noisy version of the nuclear-perinuclear localization-marker simulation, known as simulated dataset #3 (Supplementary Table 3). The same four group-specific expression and localization patterns from simulated dataset #1 were preserved, but sparse noise molecules were added to non-active gene channels. Noise was added only to gene-group combinations in a non-cell-type-specific manner (e.g., Gene 2 transcripts added to Group 1 cells). For these non-expressing gene-group pairs, we added zero to three randomly sampled transcript coordinates per cell, drawn uniformly from anywhere within the cell boundary (radius = 75 pixels). For example, in Groups 2 and 3, where only gene 2 is truly expressed, we added a small number of noise transcripts for gene 1 and gene 3. Similarly, in Group 1, we added noise for gene 2 and gene 3, and in Group 4, we added noise for gene 1 and gene 2. This dataset was used to evaluate whether lowly detected non-cell-type-specific molecules could affect image-derived features and whether feature weighting could mitigate this effect. Genes not active in a given group had zero molecules and were represented by blank black image channels after rasterization. The final simulated dataset consisted of transcript-level x, y coordinates annotated with gene, cell, and group identity, along with the derived traditional cell-by-gene count matrix for downstream evaluation and comparison.

#### 4.4.4 Nuclear-perinuclear continuum simulation dataset

To evaluate whether STARIT could capture continuous variation in subcellular transcript localization, we generated a nuclear-perinuclear continuum simulation dataset, known as simulated dataset #4 (Supplementary Table 3). This dataset contained 80 cells. Each cell expressed only gene 2, while gene1 and gene 3 were absent and represented by blank image channels. Each cell contained 80 gene 2 molecules.

For each cell, we assigned a continuum value ranging from 0 to 1, to dictate the ratio of nuclear to perinuclear transcripts of a gene. Based on this value, the 80 molecules were proportionally allocated between the nucleus and the surrounding perinuclear region. To mimic a realistic cellular geometry, molecules assigned to the nucleus were distributed uniformly by area within a circular nuclear disk, while perinuclear molecules were distributed uniformly by area within a surrounding annulus. Spatial coordinates for both regions were generated using area-proportional random sampling in polar coordinates. This design established a controlled, gradual transition from entirely nuclear to entirely perinuclear gene localization across the cell population while maintaining a constant total molecule count. The final simulated dataset consisted of transcript-level x, y coordinates annotated with gene, cell, and group identity, along with the derived traditional cell-by-gene count matrix for downstream evaluation and comparison.

#### 4.4.5 Reduced molecule-count localization-marker simulation datasets

To approximate partial transcript capture associated with thin tissue sections, we generated reduced molecule-count localization-marker simulation datasets. These simulations modeled partial capture by reducing the number of detected molecules in a subset of cells while preserving the same underlying localization patterns.

Each reduced-count dataset contained four groups of 20 cells. Group 1 expressed only gene 1 throughout the full cell. Group 2 expressed only gene 2 in the perinuclear annulus. Group 3 expressed only gene 2 in the nuclear region. Group 4 expressed only gene 3 throughout the full cell. Within each group, the first 10 cells were high-count cells with 80 molecules of the active gene. The remaining 10 cells were low-count cells with fewer molecules of the same active gene.

Three reduced-count variants were generated: 80/40 (simulated dataset #5), 80/20 (simulated dataset #6), and 80/10 (simulated dataset #7) (Supplementary Table 3). In the 80/40 dataset, the low-count cells contained 40 molecules. In the 80/20 dataset, the low-count cells contained 20 molecules. In the 80/10 dataset, the low-count cells contained 10 molecules. These variants correspond to progressively more severe reductions in molecular sampling while maintaining the same gene identity and localization-marker structure.

Molecular x,y coordinates were rounded to the nearest integer and rechecked to ensure that the rounded coordinates remained inside the intended sampling region. Genes not active in a given group had zero molecules and were represented by blank black image channels after rasterization. The final simulated dataset consisted of transcript-level x, y coordinates annotated with gene, cell, and group identity, along with the derived traditional cell-by-gene count matrix for downstream evaluation and comparison.

### 4.5 Applying STARIT to simulated imSRT datasets

STARIT was applied to each simulated dataset to generate one rasterized image per cell-gene pair with a pixel resolution size (dx) of 1 and a Gaussian kernel (blur) of 1. For datasets with three genes, three images were produced for each cell. If a cell had zero molecules for a given gene, a blank black 224×224 image was generated for that gene. For ResNet101-based simulated analyses, the three gene-specific embeddings were concatenated to form an 80 x 6,144 cell-by-feature matrix. For ResNet18, ResNet50, ResNet152, CTransPath, and UNI2-h comparisons, the same image preprocessing and concatenation strategy was used, with feature dimensions determined by the corresponding model architecture and reported in Supplementary Table 1.

### 4.6 Downstream analysis of simulated datasets

For STARIT-based analyses of the main simulated datasets, PCA was first applied to the cell-by-feature matrix. Unless otherwise specified, the top 5 principal components were retained. UMAP was then applied to the PCA representation for visualization. For the main ResNet101 nuclear-perinuclear simulation, UMAP used 30 nearest neighbors, a minimum distance of 0.1, and random state 42. For most additional simulated STARIT analyses, UMAP used 30 nearest neighbors, a minimum distance of 0.2, and random state 42, unless otherwise indicated in Supplementary Table 1, keeping all other parameters at their defaults.

Graph-based clustering was performed by constructing a k-nearest neighbors (kNN) graph from the PCA representation using cosine distance, connectivity mode, and include self to true. Louvain clustering was then applied to the resulting graph. The number of neighbors and Louvain resolution were selected for each analysis as reported in Supplementary Table 1. For the main ResNet101 nuclear-perinuclear simulation, 18 neighbors were used. For feature-weighted noisy simulated analyses, PCA retained 5 PCs, UMAP used 10 nearest neighbors and a minimum distance of 0.5, and Louvain clustering was applied to a kNN graph constructed with the number of neighbors specified in Supplementary Table 1.

To compare STARIT-derived clusters with ground-truth simulated groups, we computed the adjusted rand index (ARI) and a Jaccard similarity matrix. ARI was used to summarize agreement between the complete inferred clustering and ground-truth annotations while correcting for chance agreement. Jaccard similarity was computed between each ground-truth group and each inferred cluster. When the number of inferred clusters differed from the number of ground-truth groups, mean on-diagonal Jaccard similarity was computed from the best one-to-one matching between groups and clusters, such as the case for certain models with the left-right polarized simulated dataset.

For conventional gene-count analysis of simulated datasets, PCA was applied to the cell-by-gene count matrix rather than the STARIT feature matrix. Unless otherwise specified, the top 3 PCs were retained. UMAP and Louvain clustering were then performed using the parameters reported in Supplementary Table 1. Cluster agreement with ground truth was quantified using the same ARI and Jaccard similarity approaches used for STARIT-derived clusters, where again when the number of inferred clusters differed from the number of ground-truth groups, mean on-diagonal Jaccard similarity was computed from the best one-to-one matching between groups and clusters.

For selected analyses, extracted image features were visualized using heatmaps. Feature values were standardized across cells by z-score normalization. Hierarchical clustering was performed using Ward’s linkage and Euclidean distance.

### 4.7 Differential feature testing and differential expression analyses

To identify STARIT-derived features that differed between selected clusters, we applied two-sided Wilcoxon rank-sum tests to extracted image features. P-values were adjusted for multiple testing using Bonferroni correction. For the simulated datasets, differential testing was performed on STARIT-derived image features to identify features associated with group-specific gene images and localization patterns.

For the osmFISH analysis, differential testing was performed between selected STARIT-derived clusters using both STARIT image features and conventional gene counts. For STARIT feature testing, Wilcoxon rank-sum tests were applied to the image-derived feature matrix. For conventional expression testing, the same tests were applied to the cell-by-gene count matrix. For the bacterial-MERFISH analysis, differential testing was also performed between the selected STARIT-derived clusters using STARIT-derived image features.

Genes or features with Bonferroni-corrected p-values below the specified threshold of 0.001 were considered significant.

### 4.8 osmFISH Mouse Cortex Data

The osmFISH mouse somatosensory cortex dataset was obtained from the original publication by Codeluppi, *et al*. [13]. Cell regions annotated as “Excluded” in the original dataset were removed. We also removed six additional cells with no detected molecules for any of the 33 assayed genes (cells 7825, 1524, 3085, 5619, 7823, 4323). After filtering, the dataset contained 4,833 cells, 33 genes, 11 curated anatomical regions, and 31 curated cell-type annotations. For each cell, subcellular transcript coordinates were centered at the cell origin. STARIT was then applied to all 4,833 cells and all 33 genes using a pixel resolution size of 1 (dx = 1) and a Gaussian kernel of 1 (blur = 1). This generated 159,489 gene-specific rasterized images. For genes with zero detected molecules in a given cell, a blank black 224×224 image was generated.

### 4.9 Downstream STARIT analysis of osmFISH data

Each STARIT image was converted to RGB, resized to 224×224 pixels with aspect-ratio preservation and padding, and normalized with ImageNet preprocessing. A pretrained ResNet101 with the final fully connected layer removed was used to extract a 2,048-dimensional feature embedding for each gene image. Because there are thirty-three gene markers, 33 corresponding images were produced. The 2048-dimensional ResNet features extracted from each gene-specific image were concatenated to form a 67,584-dimensional feature vector per cell (33 genes x 2,048 features). Feature vectors from all 4,833 cells were then stacked into a 4,833 x 67,584 matrix, which served as the input for all downstream analyses.

For the unweighted STARIT analysis, PCA was applied to the feature matrix, and the top 30 PCs were retained. UMAP was applied using cosine distance, 50 nearest neighbors, a minimum distance of 0.15, and random state 42. For clustering, a kNN graph was constructed in PCA space using cosine distance, connectivity mode, include self to true, and 5 nearest neighbors. Louvain clustering was applied with the parameters reported in Supplementary Table 1. To compare with the 31 curated osmFISH cell types, hierarchical agglomerative clustering was performed on Louvain community centroids in PCA space to obtain 31 meta-clusters. Agreement between inferred clusters and curated cell-type annotations was quantified using ARI and Jaccard similarity.

For the feature-weighted osmFISH STARIT analysis, the original osmFISH cell-by-gene count matrix was min–max scaled per gene across cells. During feature extraction for each cell and gene, the corresponding 2,048-dimensional ResNet101 embedding was multiplied by the scaled count value for that gene in that cell. Weighted gene-specific embeddings were concatenated to generate a 4,833 x 67,584 weighted cell-by-feature matrix. PCA, UMAP, Louvain clustering, and agglomerative clustering were then performed as described above, except that the kNN graph for Louvain clustering used 6 nearest neighbors, as reported in Supplementary Table 1. Agreement between inferred clusters and curated cell-type annotations was once again quantified using ARI and Jaccard similarity.

### 4.10 Conventional gene-count analysis of osmFISH data

Using the original osmFISH cell-by-gene count matrix, gene counts were filtered identically and min–max scaled per gene across cells (4,833 cells x 33 genes). We first performed PCA and retained the top 30 PCs. These PCs were then used as input to UMAP using, nearest neighbors of 50, a minimum distance of 0.15, and random state 42. A kNN graph was constructed from the PCA representation using 6 nearest neighbors, cosine distance, include self to true, and connectivity mode, keeping all other settings at their defaults. Louvain clustering was performed with a resolution of 1.2 and random state 42. Hierarchical agglomerative clustering was then applied to Louvain community centroids in PCA space to obtain 31 meta-clusters. Agreement with curated osmFISH cell-type labels was quantified using ARI and Jaccard similarity.

### 4.11 Bacteria-MERFISH data

The 1000-fold volumetric expansion bacteria-MERFISH dataset was obtained from the original publication Sarfatis *et al.* [14]. For the 1000-fold bacterial-MERFISH dataset, we filtered for bacterial cells with more than 250 total mRNA molecules detected, resulting in 463 cells. For each retained cell, molecular coordinates were subsetted to those corresponding to the fliC–fliX operon. Subcellular positional coordinates were centered at each cell’s origin prior to rasterization.

STARIT was applied to these centered fliC-fliX molecular coordinates, with a pixel resolution size of 1 (dx=1) and a Gaussian kernel of 1 (blur=1), producing one rasterized STARIT image per cell. A total of 463 operon image representations of fliC-fliX were generated.

### 4.12 Downstream analysis of unstandardized bacterial-MERFISH data

For the unstandardized bacterial-MERFISH analysis, each STARIT image was converted to RGB, resized to 224×224 pixels with aspect-ratio preservation and padding, and normalized using ImageNet preprocessing. A pretrained ResNet101 with the final fully connected layer removed was used to extract a 2,048-dimensional feature embedding for each operon image to form a 2,048-dimensional feature vector per cell (1 operon x 2,048 features). Feature vectors from all 463 cells were then stacked into a 463 x 2,048 matrix.

PCA was applied to the feature matrix and the top 30 PCs were retained. UMAP was performed using 15 nearest neighbors, a minimum distance of 0.1, and random state 42. For clustering, a kNN graph was constructed in PCA space using cosine distance, connectivity mode, include self to true, and 100 nearest neighbors. Louvain clustering was then applied using random state 42, keeping all other settings at their defaults. Cluster-specific variation in fliC–fliX expression was visualized using PCA embeddings colored by raw and log-transformed counts.

### 4.13 Rotation standardization of bacterial-MERFISH cells

To reduce clustering driven by whole-cell orientation, we generated a rotation-standardized bacterial-MERFISH data representation. For each bacterium, fliC-fliX molecular coordinates were centered translating the origin to the global centroid, calculated from all detected operon molecules within the cell.

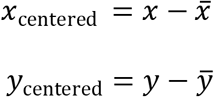

PCA was then applied to the centered fliC-fliX coordinates to identify the first principal component eigenvector, ***v*** = (*v*_1_, *v*_2_)*^T^*, corresponding to the major axis of spatial variation. The orientation angle between this first principal component and the x-axis was computed as *θ*, using the four-quadrant inverse tangent:

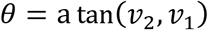

A two-dimensional counterclockwise rotation transformation was then applied to align each cell’s major axis with the x-axis prior to rasterization, using the angle *θ*:

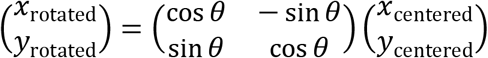

This standardization was used to reduce variation associated with arbitrary bacterial orientation.

### 4.14 Downstream analysis of rotation-standardized bacterial-MERFISH data

STARIT was applied to the rotation-standardized fliC–fliX coordinates to generate one rasterized operon image per cell. Images were converted to RGB, resized to 224×224 pixels with aspect-ratio preservation and padding, and normalized using ImageNet preprocessing.

For the standardized bacterial-MERFISH analysis, UNI2-h was used for feature extraction, producing a 463 x 1,536 cell-by-feature matrix. PCA was applied and the top 30 PCs were retained. UMAP was performed using 15 nearest neighbors, a minimum distance of 0.1, and random state 42. A kNN graph was constructed in PCA space using cosine distance, connectivity mode, include self to true, and 100 nearest neighbors. Louvain clustering was then applied using random state 42, keeping all other settings at their defaults.

### 4.15 Interpretable image-feature analysis of bacterial-MERFISH clusters

To characterize the STARIT-derived bacterial-MERFISH clusters using interpretable measurements, we extracted supervised morphological and intensity metrics from each cell. Metrics included spatial spread of fliC-fliX gene expression in the x-direction, total sum pixel intensity, total normalized expression of fliC-fliX, cell length, and total operon expression. In the absence of experimentally provided cell segmentation from the Bacterial-MERFISH data, we defined a proxy cell segmentation mask for each cell using the molecular coordinate data provided from Sarfatis *et al*. Specifically, we used STARIT to generate a cell segmentation image per cell using all detected molecules assigned to all operons from their spatial coordinates. The resulting image was converted to grayscale and thresholded (values greater than 0) to generate a binary cell segmentation mask. This proxy cell mask was used to approximate the cell region over which intensity-based metrics were quantified, such as total sum intensity, summing up all the pixel intensities from each fliC-fliX image that are inside the cell mask. Calculations for spatial spread of fliC-fliX gene expression in the x-direction and cell length per cell were derived from the actual *x,y* molecular coordinate information rather than the images themselves. A full list defining each of these metrics and how they are calculated are in Supplementary Table 2.

For each metric, pairwise differences between clusters were evaluated using two-sided Wilcoxon rank-sum tests. P-values were adjusted for multiple testing using Bonferroni correction. These supervised metrics were used to summarize interpretable differences between bacterial clusters, while recognizing that STARIT-derived cluster structure may reflect multivariate combinations of spatial and intensity features rather than a single supervised metric.

Spearman correlations were also computed between extracted image features and each supervised metric.

### 4.16 Computational Specifications

All STARIT rasterization and downstream feature extraction runtimes reported were measured in series, without any parallelization across cells or genes, on a Mac Studio (2023) equipped with an Apple M2 Ultra system-on-chip (24-core CPU: 16 performance cores and 8 efficiency cores; 60-core GPU), 192 GB of unified memory, running macOS Sequoia 15.6.1.

## Data availability

All data analyzed with STARIT are publicly available. The simulated datasets are available at https://github.com/JEFworks-Lab/STARIT/tree/main/data. The osmFISH datasets are available at https://linnarssonlab.org/osmFISH/availability/. The Bacterial-MERFISH dataset is available at https://datadryad.org/dataset/doi:10.5061/dryad.n5tb2rc4d.

STARIT is available as an open-source Python toolkit, with additional documentation and tutorials available at https://github.com/JEFworks-Lab/STARIT.

## Funding

This work was supported by the National Science Foundation [2047611] and the National Institute of General Medical Sciences of the National Institutes of Health [R35-GM142889].

## Competing interests

The authors have declared that no competing interests exist.

## Supplementary Material

### Supplementary Tables

**Table 1.** Summary of datasets, analysis configurations, and clustering performance metrics. This table summarizes the STARIT and baseline analysis configurations applied to each dataset. For each dataset and feature extraction strategy, the table reports the resulting feature matrix size, whether feature weighting was applied, the clustering workflow and parameter settings, the number of clusters identified, and clustering performance metrics where ground-truth labels were available. Performance was evaluated using Adjusted Rand Index and mean on-diagonal Jaccard similarity. Entries labeled “N/A” indicate analyses where ground-truth labels or direct cluster-to-label comparisons were not applicable.

| Column Name | Description |
| --- | --- |
| Dataset ID | Dataset identifier corresponding to the dataset ID in Table 1. |
| Dataset Name | Name of the dataset analyzed. |
| Feature Extractor | Feature representation or model used to extract features from STARIT image tensors, such as ResNet101, UNI2-h, CTransPath, or N/A for gene-counts based analysis. |
| Feature Matrix | Dimensions of the feature matrix used for downstream analysis, reported as number of cells by number of features. |
| Feature Weighting | Indicates whether feature weighting was applied before dimensionality reduction and clustering. |
| Number of Clusters | Number of clusters identified by the clustering workflow. |
| Clustering Approach | Dimensionality reduction and clustering workflow used, such as PCA, UMAP, Louvain clustering, and, where applicable, agglomerative clustering. |
| PCA Parameters | Principal component analysis parameters used for dimensionality reduction, including the number of components and random seed. |
| UMAP Parameters | UMAP embedding parameters, including number of neighbors, minimum distance, and random seed. |
| kNN Graph Parameters | Parameters used to construct the k-nearest-neighbor graph for clustering, including neighbor number, distance metric, graph mode, and whether self-neighbors were included. |
| Louvain Clustering Parameters | Louvain clustering parameters, including random seed and resolution when specified. |
| Adjusted Rand Index (ARI) | Adjusted Rand Index comparing identified clusters to known labels, where available. Higher values indicate stronger agreement between clustering results and reference labels. |
| Mean on-diagonal Jaccard Similarity | Mean Jaccard similarity between matched clusters and reference labels along the diagonal after cluster-label matching. Higher values indicate stronger overlap. |

**Table 2.** Supervised metrics for standardized Bacterial-MERFISH *E. coli* data. This table reports the supervised spatial, morphological, and transcriptomic metrics evaluated across the identified clusters in the standardized Bacterial-MERFISH *E. coli* dataset.

| Column ID | Description |
| --- | --- |
| Metric Name | Name of supervised metric. |
| Description | Definition of metric. |
| Calculation | Description of how metric was calculated. |

**Table 3.** Overview of datasets. This table summarizes all datasets used in this study, including simulated imSRT datasets and real spatial transcriptomics datasets. For each dataset, the table provides a dataset identifier, dataset name, brief description, number of cells, number of genes or assayed targets, and number of annotated cell types or cell states where applicable.

| Column Name | Description |
| --- | --- |
| ID | Numeric identifier assigned to each dataset. |
| Name | Dataset name used throughout the analysis. |
| Description | Brief description of the dataset, including the biological or simulation context and the spatial localization pattern being evaluated. |
| Cells | Number of cells included in the dataset after filtering or simulation. |
| Genes | Number of genes, targets, or transcript channels included in the dataset. |
| Cell-Types | Number of known cell types, simulated cell states, or annotated groups used for comparison. “N/A” indicates that discrete cell-type labels were not applicable. |

### Supplementary Figures

**Supplementary Figure 1.**
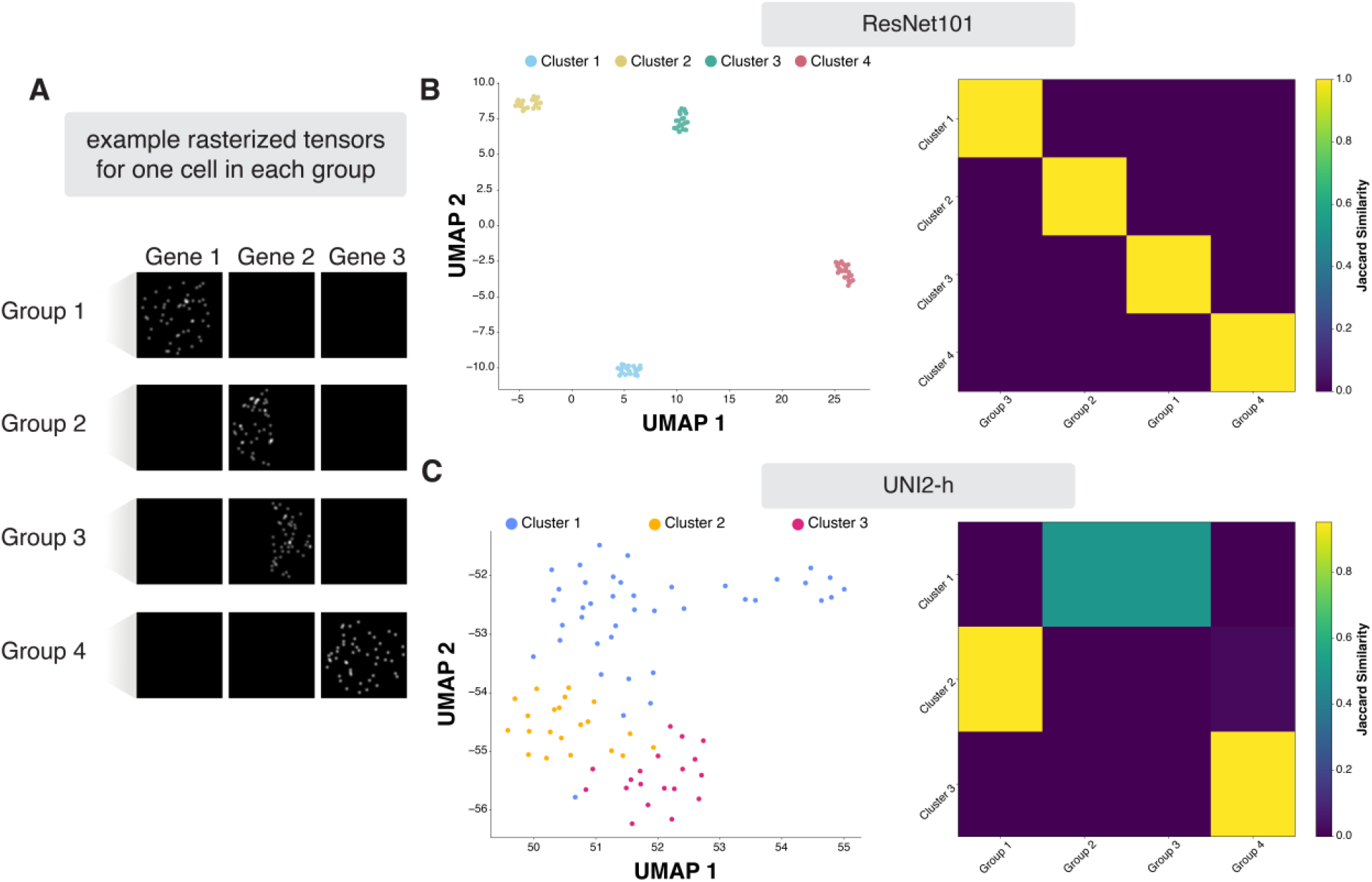
Comparison of STARIT-derived ResNet101 features versus STARIT-derived UNI2-h features for left-right polarized localization simulation imSRT dataset. (A) Representative STARIT rasterized image tensors for one cell from each of the four simulated groups. (B) Analysis results of STARIT tensors using ResNet101 feature extraction. Left: UMAP visualization based on STARIT output, embedding colored by Louvain cluster assignment. Right: Jaccard similarity heatmap between ground-truth and STARIT-based Louvain clusters. (C) Analysis results of STARIT tensors using UNI2-h feature extraction. Left: UMAP visualization based on STARIT output, embedding colored by Louvain cluster assignment. Right: Jaccard similarity heatmap between ground-truth and STARIT-based Louvain clusters.

**Supplementary Figure 2.**
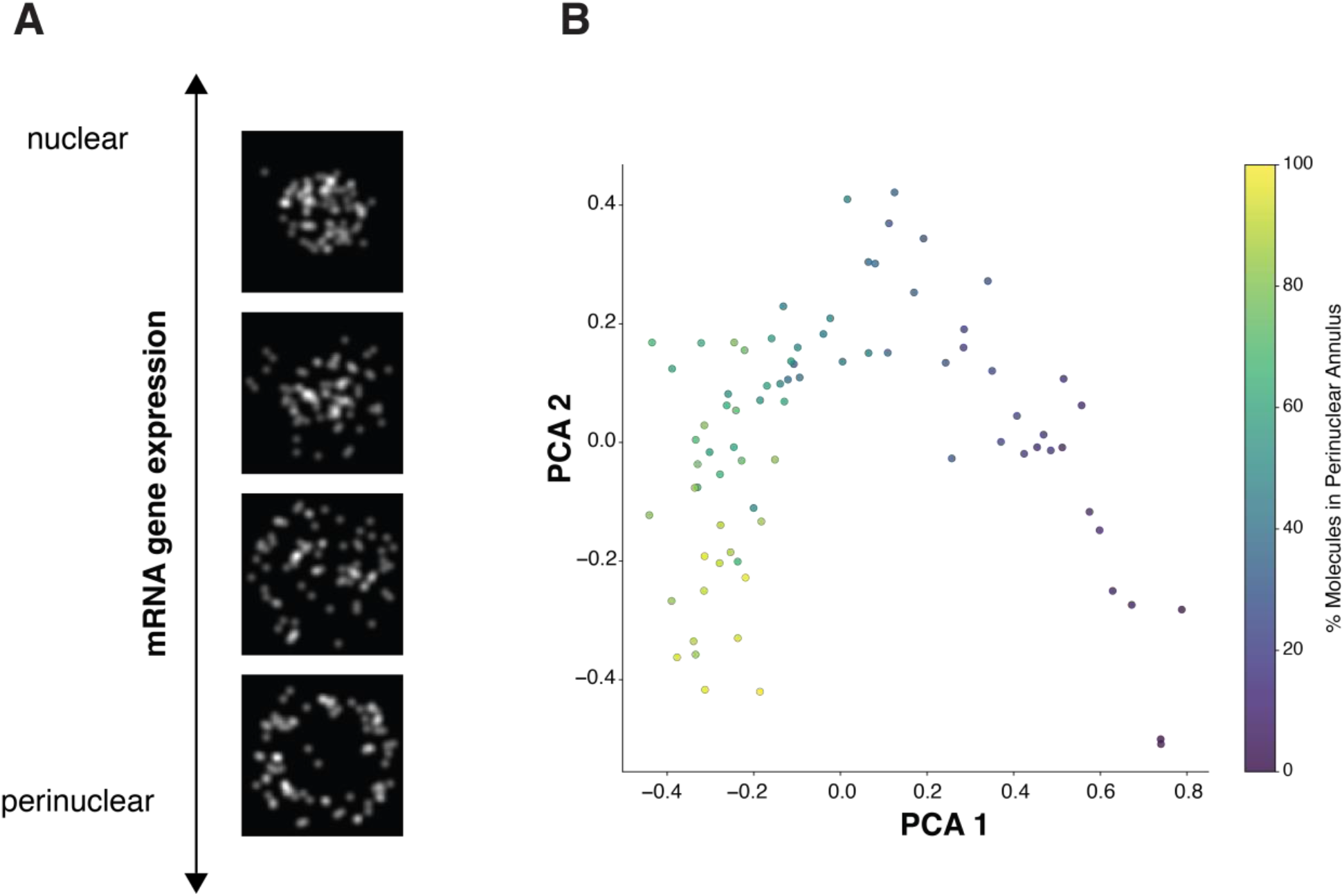
STARIT-derived ResNet101 features for the nuclear-perinuclear continuum simulation dataset. (A) Representative cell examples from the nuclear-perinuclear continuum simulation dataset, illustrating the spatial transition of mRNA transcript localization from the nucleus (top) to the outer perinuclear annulus (bottom). (B) Principal Component Analysis (PCA) projection of extracted ResNet101 features across simulated cells, colored by the percentage of molecules localized to the perinuclear annulus.

**Supplementary Figure 3.**
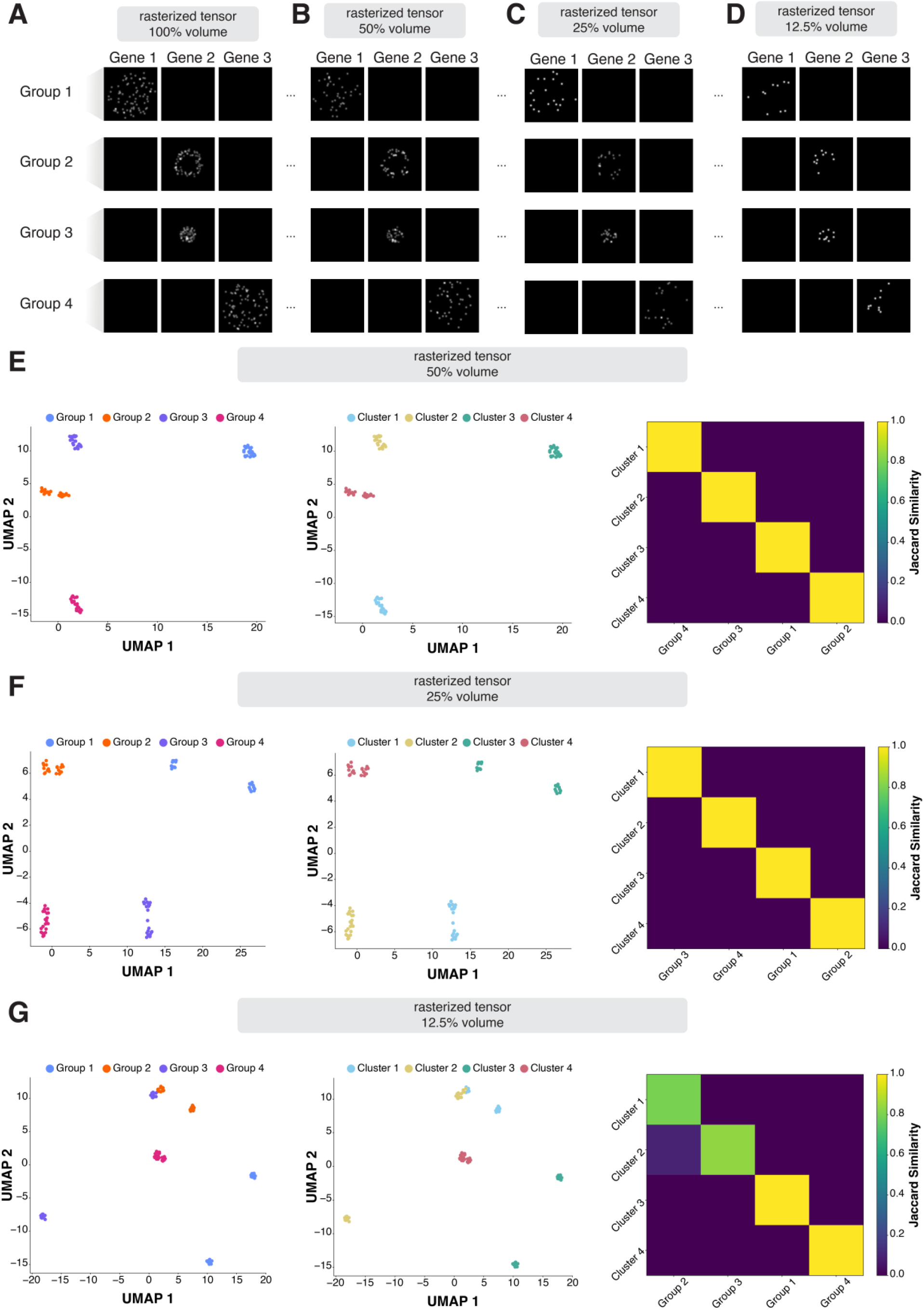
STARIT-derived ResNet101 features for simulated cut-volume molecule count localization datasets. (A-D) Representative STARIT rasterized image tensors for one cell from each simulated group at (A) 100%, (B) 50%, (C) 25%, and (D) 12.5% retained cell volume. Within each panel, columns correspond to Gene 1, Gene 2, and Gene 3, and rows correspond to the four simulated groups. (E-G) Clustering analyses for simulations with (E) 50%, (F) 25%, and (G) 12.5% retained cell volume. (E) Left: UMAP visualization based on ground-truth, with embedding colored by group. Middle: UMAP visualization based on STARIT output, with embedding colored by Louvain cluster assignment. Right: Jaccard similarity heatmap between ground-truth and STARIT-based Louvain clusters. (F) Left: UMAP visualization based on ground-truth, with embedding colored by group. Middle: UMAP visualization based on STARIT output, with embedding colored by Louvain cluster assignment. Right: Jaccard similarity heatmap between ground-truth and STARIT-based Louvain clusters. (G) Left: UMAP visualization based on ground-truth, with embedding colored by group. Middle: UMAP visualization based on STARIT output, with embedding colored by Louvain cluster assignment. Right: Jaccard similarity heatmap between ground-truth and STARIT-based Louvain clusters.

**Supplementary Figure 4.**
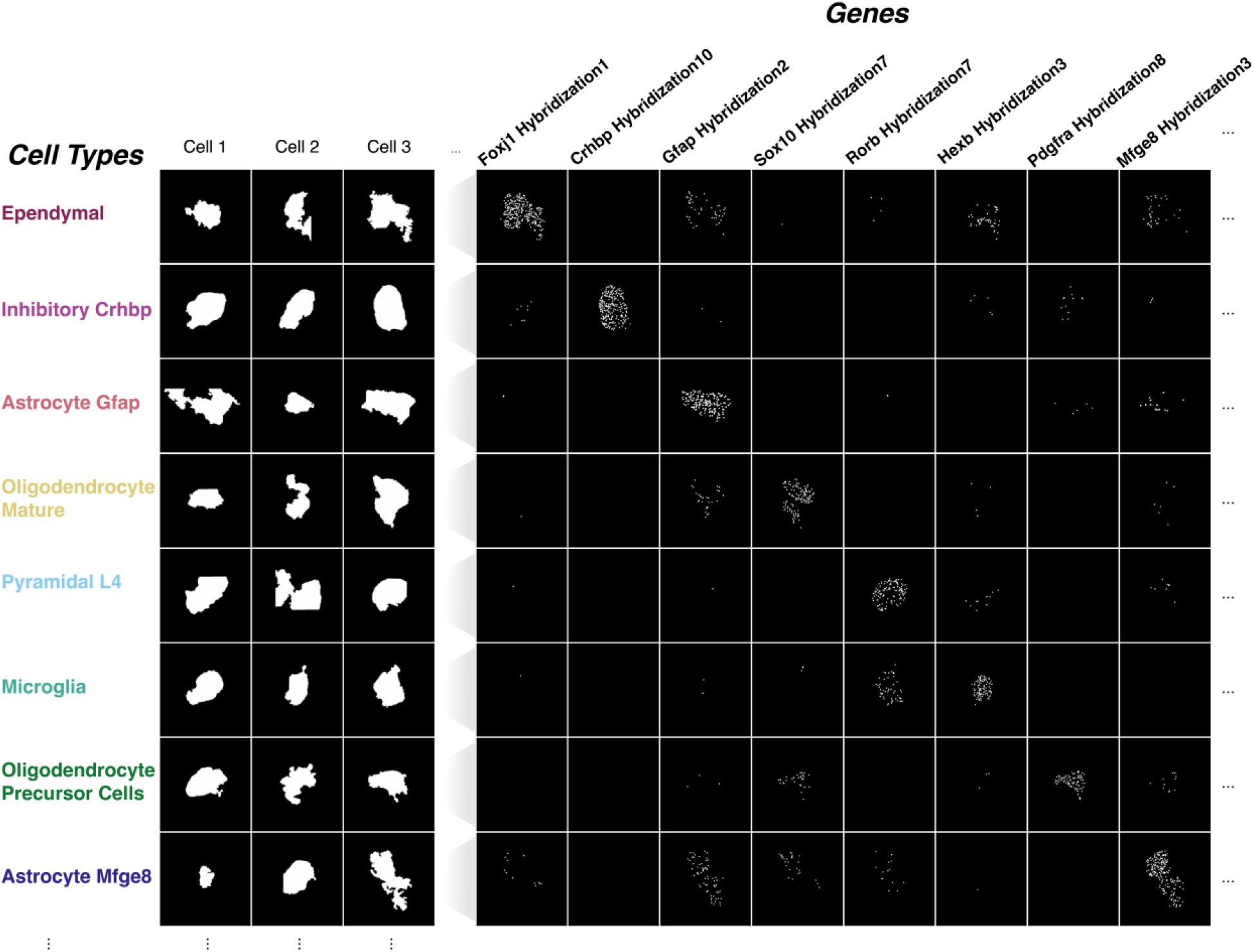
Example cell segmentation and gene-specific STARIT rasterized images from osmFISH mouse cortex data cell types. Representative cell segmentation and corresponding STARIT-generated gene rasterizations for selected osmFISH-defined cell types and genes. Left: For each of eight example cell types (Ependymal, Inhibitory Crhbp, Astrocyte Gfap, Oligodendrocyte Mature, Pyramidal L4, Microglia, Oligodendrocyte Precursor Cells, and Astrocyte Mfge8), we show three individual cell segmentation masks. Right: For one of the cells of each cell type, we display STARIT gene tensor rasterizations for a subset of the 33 assayed genes, where each panel shows the spatial distribution of transcripts within the cell segmentation mask boundary.

**Supplementary Figure 5.**
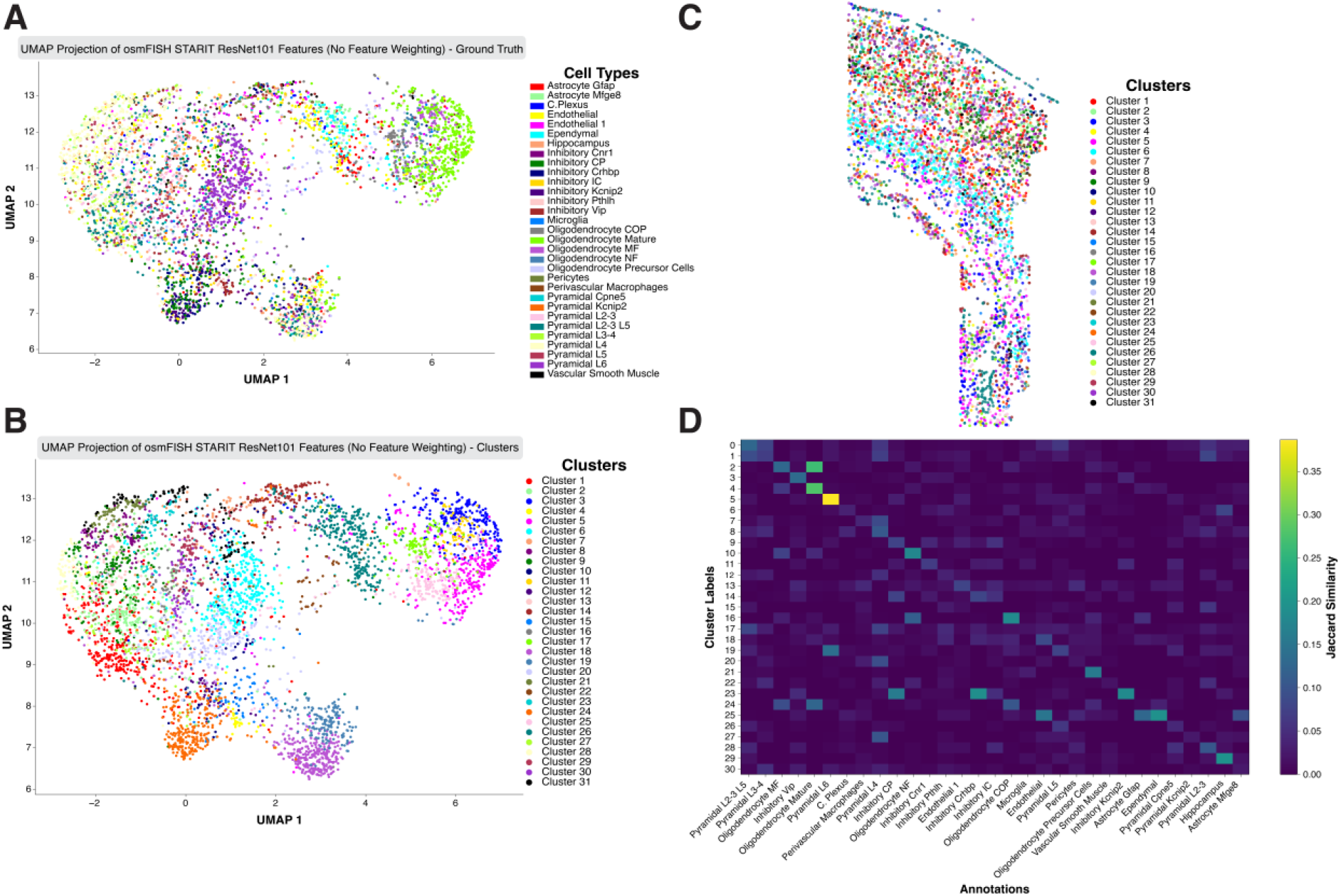
STARIT feature embedding and clustering of osmFISH mouse cortex cells and comparison to previously annotated cell types, without feature weighting. (A) UMAP visualization of STARIT-derived features for 4,833 osmFISH cells from the mouse cortex, colored by the 31 original cell-type annotations reported by Codeluppi *et al.* (2018). Each point represents a single cell. (B) UMAP embedding of the same STARIT features colored by 31 STARIT-derived Louvain clusters and subsequent agglomerative clustering. Each point represents a single cell. (C) STARIT-derived Louvain clustering of osmFISH mouse somatosensory cortex dataset in tissue space. (D) Jaccard similarity matrix comparing STARIT-derived Louvain clusters (rows) to ground-truth cell-type annotations (columns).

**Supplementary Figure 6.**
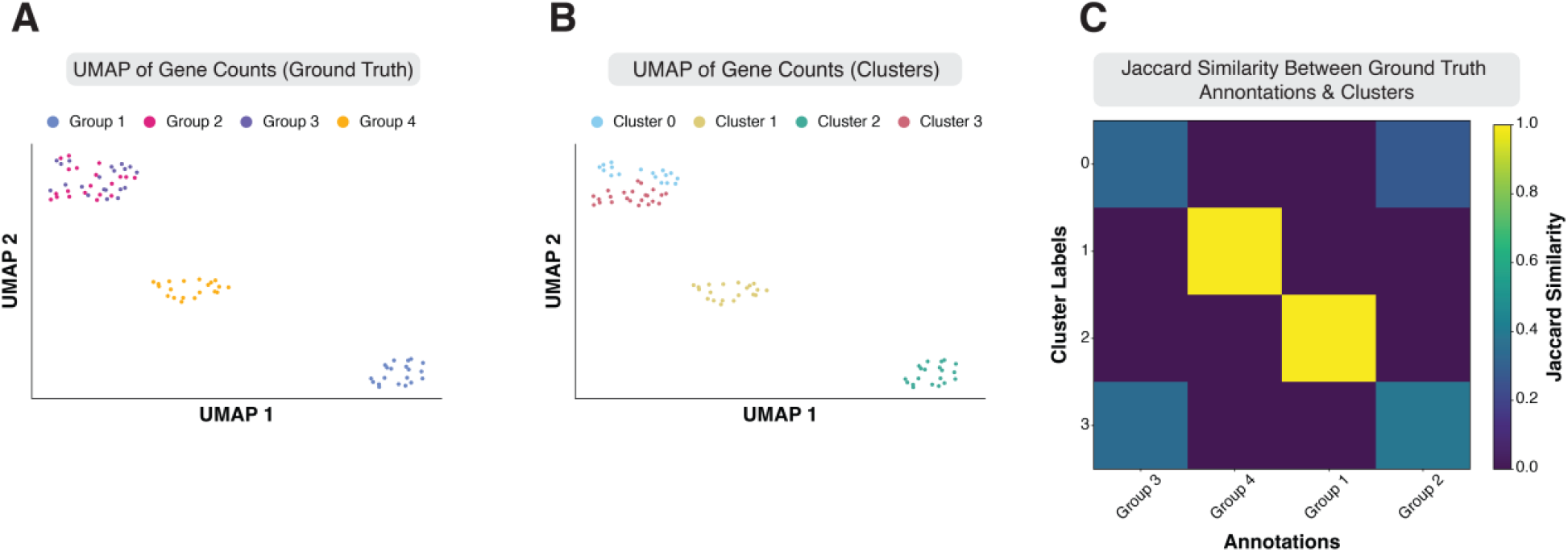
Effect of misallocated mRNA molecules on conventional gene-count–based clustering on simulated noisy imSRT data. (A) UMAP embedding of PCs from the gene count matrix is shown for ground-truth simulated noisy imSRT group annotations. (B) UMAP embedding of PCs from the gene count matrix is shown for Louvain clusters. (C) Jaccard similarity heatmap compares Louvain clusters and ground-truth simulated noisy imSRT group annotations.

**Supplementary Figure 7.**
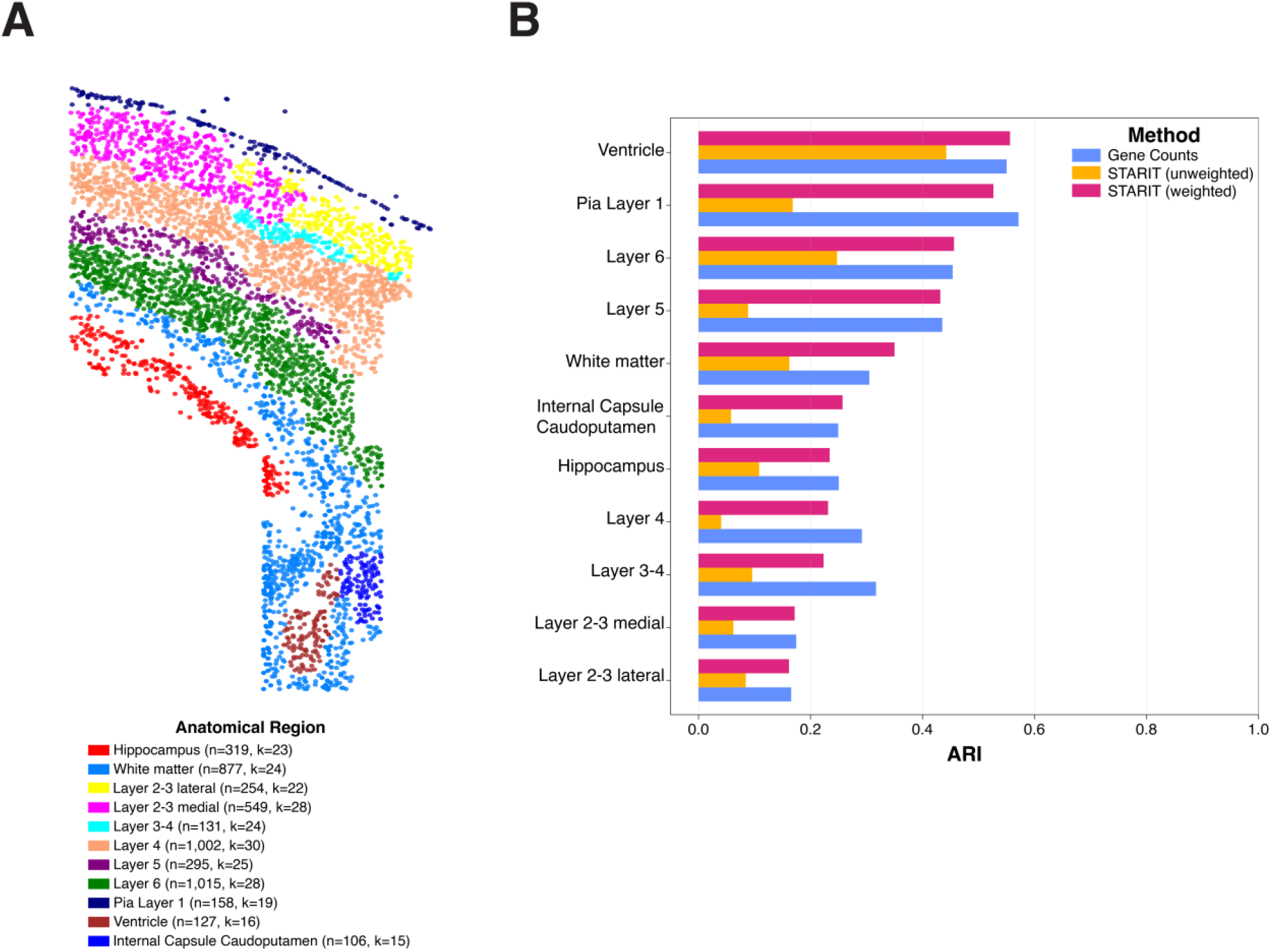
Clustering agreements of osmFISH ground-truth cell types by anatomical region. (A) Visualization of the 4,833 profiled cells in the osmFISH mouse somatosensory cortex dataset reported by Codeluppi *et al.* (2018) in tissue space, colored by the 11 annotated anatomical regions. Legend indicates the number of cells (*n*) and ground-truth annotated cell types (*k*) within each region. (B) Grouped horizontal bar chart evaluating Adjusted Rand Index (ARI) (x-axis) scores comparing ground-truth cell-type annotations with Louvain clusters derived from scaled gene counts (blue), unweighted STARIT image features (orange), and weighted STARIT image features (pink) across anatomical regions (y-axis). Regions are ordered by weighted STARIT ARI score (descending).

**Supplementary Figure 8.**
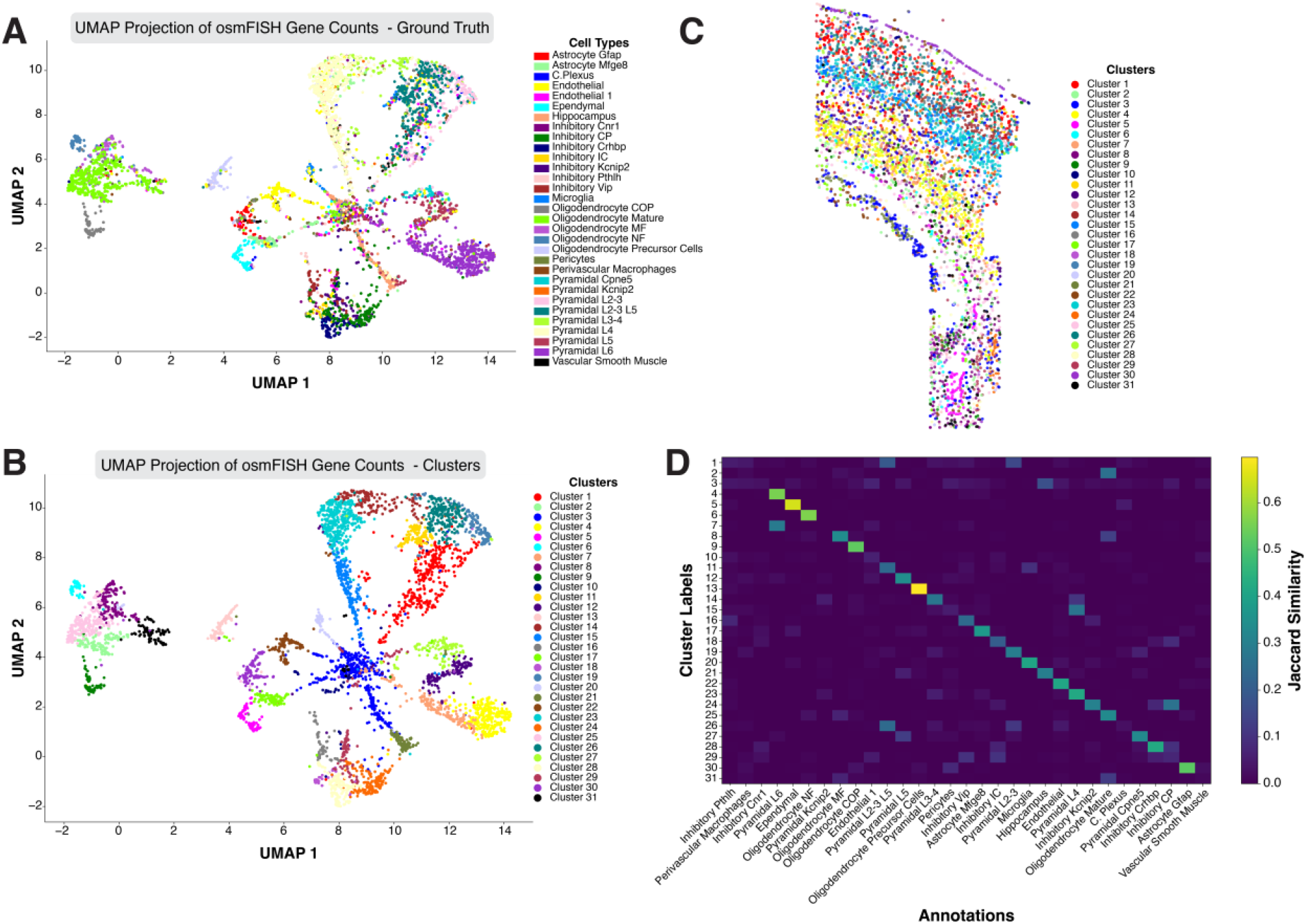
Gene count–based clustering of osmFISH mouse cortex data. (A) UMAP embedding of PC**s** from the gene count matrix of the osmFISH mouse cortex dataset, colored by the 31 previously annotated cell types from *Codeluppi et al.* (2018). (B) UMAP embedding of the same gene count–derived features colored by 31 Louvain clusters from STARIT-derived ResNet101 image features followed by agglomerative clustering. (C) Gene counts clustering of osmFISH mouse somatosensory cortex dataset in tissue space. (D) Jaccard similarity matrix comparing gene count–derived Louvain clusters (rows) to annotated cell types (columns).

**Supplementary Figure 9.**
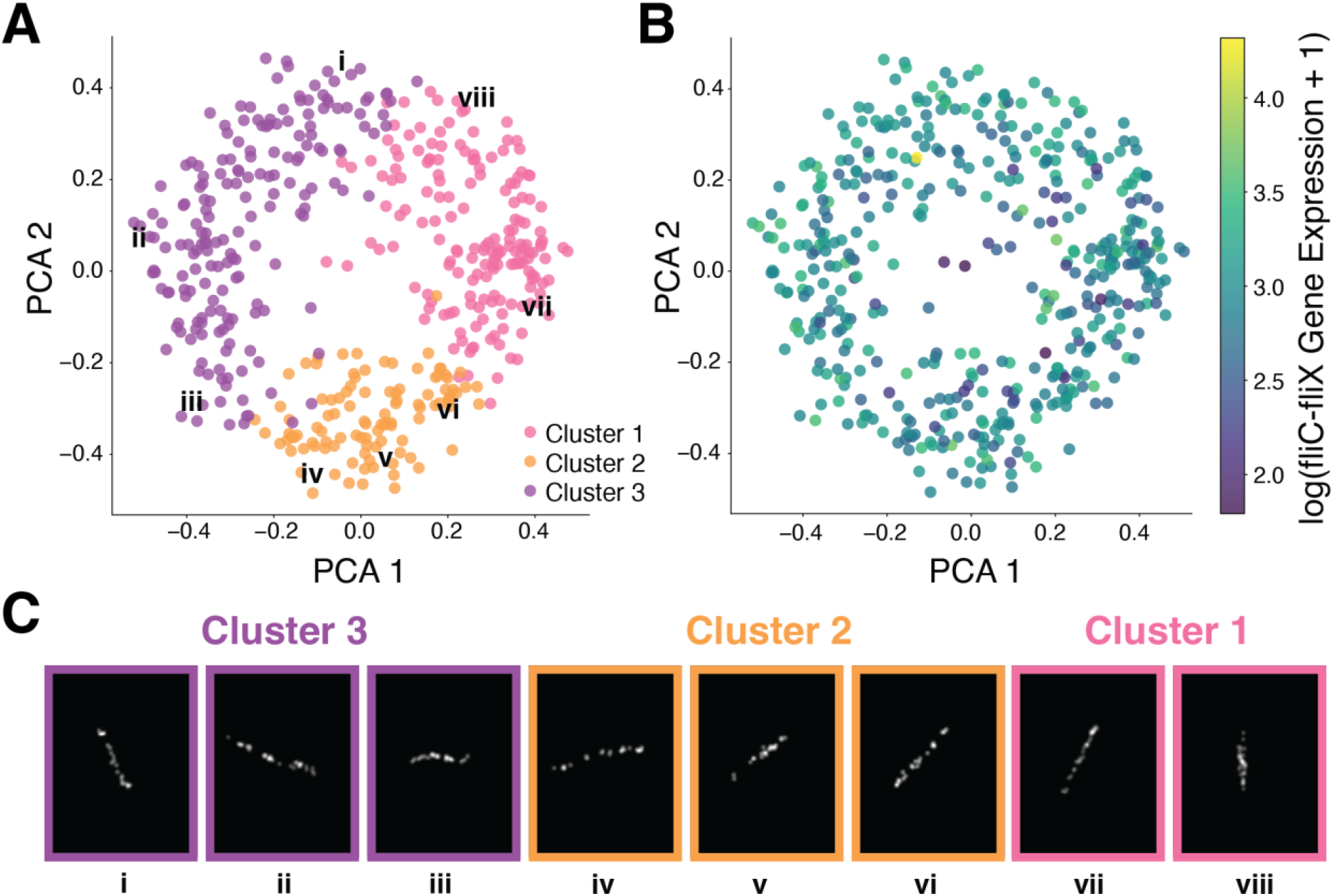
STARIT performance on unstandardized bacterial-MERFISH 1000X data. (A) Principal component analysis (PCA) of STARIT-derived features from 463 *E. coli* cells based on fliC–fliX operon localization, imaged with bacterial-MERFISH at 1000X volumetric expansion. Cells are colored by Louvain graph-based cluster assignment: Cluster 1 (pink), Cluster 2 (orange), Cluster 3 (purple). Select labeled cells (i–viii) correspond to examples shown in panel C. (B) PCA embedding of cells colored by log-transformed fliC–fliX gene expression levels. (C) Representative rasterized fliC–fliX transcript distributions for cells i–viii labeled in panel A, with frame colors indicating cluster assignment.

**Supplementary Figure 10.**
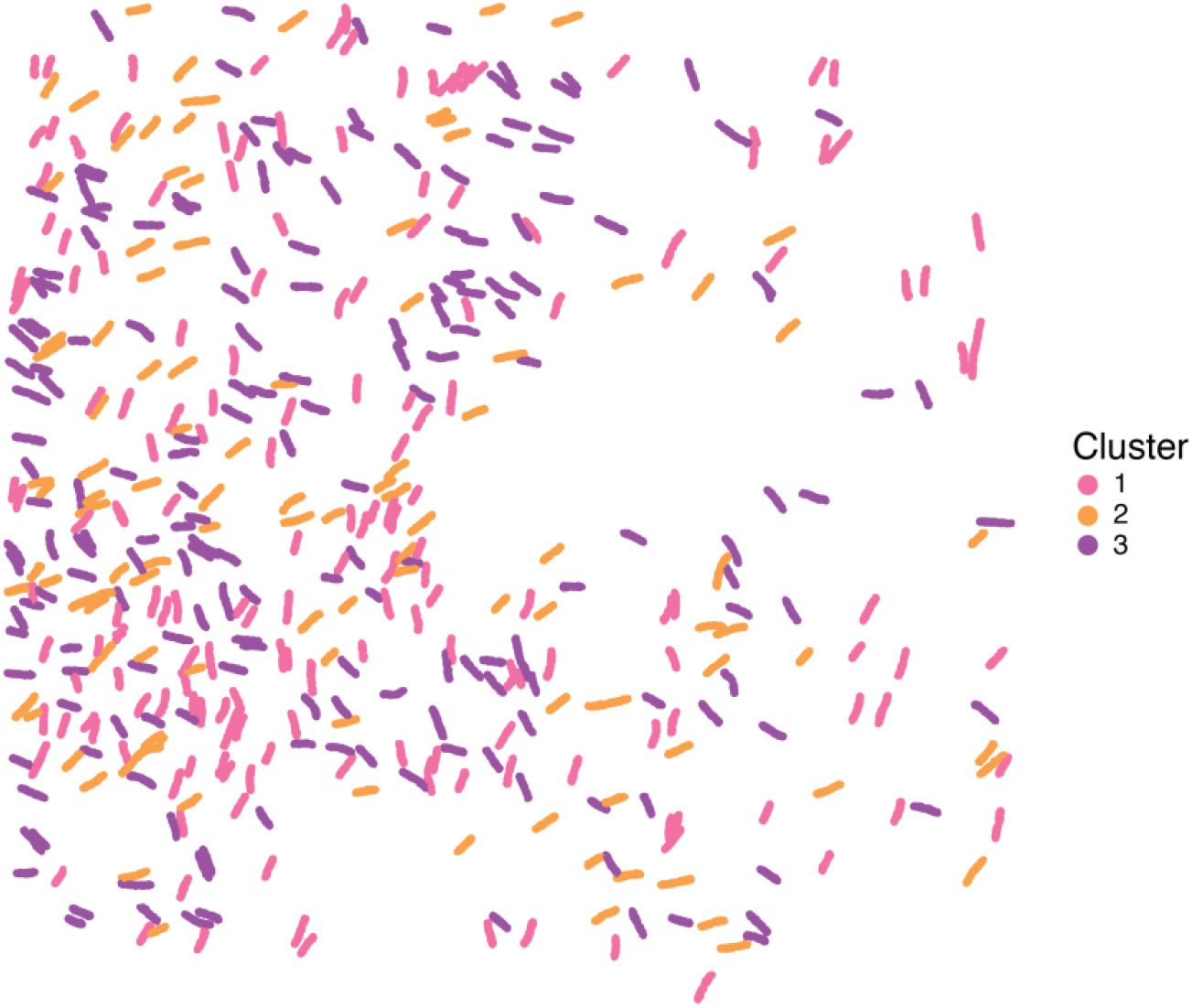
Physical-space visualization of unstandardized STARIT-identified clusters in Bacterial-MERFISH *E. coli* data. Shown are the spatial positions of 463 *E. coli* cells assayed by Bacterial-MERFISH after 1000-fold volumetric expansion, plotted in their original physical coordinates and colored by the three Louvain clusters identified from STARIT-derived ResNet101 image features.

**Supplementary Figure 11.**
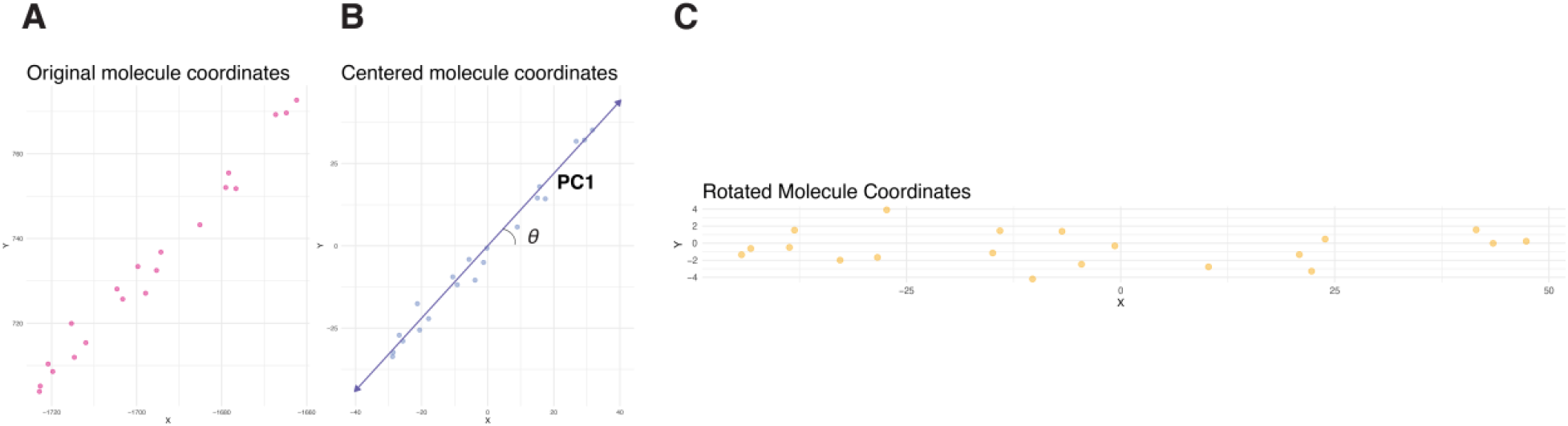
Rotational standardization of bacterial-MERFISH molecular coordinates through centering and principal axis alignment. Visualization of the coordinate-standardization procedure for one representative *E. coli* cell. (A) Original spatial coordinates of the detected fliC-fliX molecules. (B) Molecular coordinates after centering by subtraction of the coordinate centroid. The first principal component, PC1, indicates the major axis of the molecular distribution, and *θ* denotes its angle relative to the horizontal axis. (C) Centered molecular coordinates after rotation by the estimated angle to align PC1 with the horizontal axis. Each point represents one detected molecule.

**Supplementary Figure 12.**
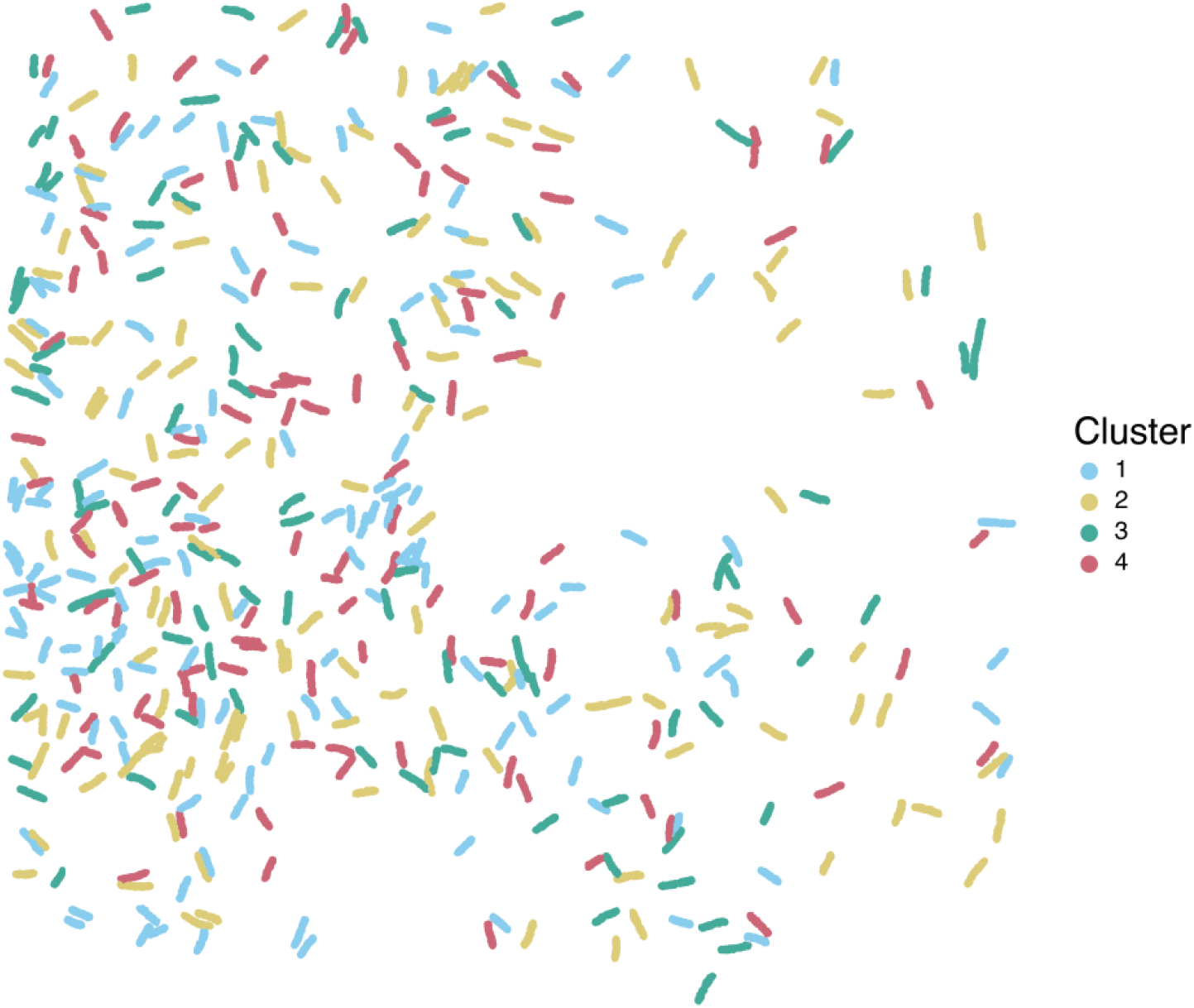
Physical-space visualization of rotationally-standardized STARIT-identified clusters in Bacterial-MERFISH *E. coli* data. Shown are the spatial positions of 463 *E. coli* cells assayed by Bacterial-MERFISH after 1000-fold volumetric expansion, plotted in their original physical coordinates and colored by the four Louvain clusters identified from **r**otationally-standardized STARIT-derived UNI2-h image features.

**Supplementary Figure 13.**
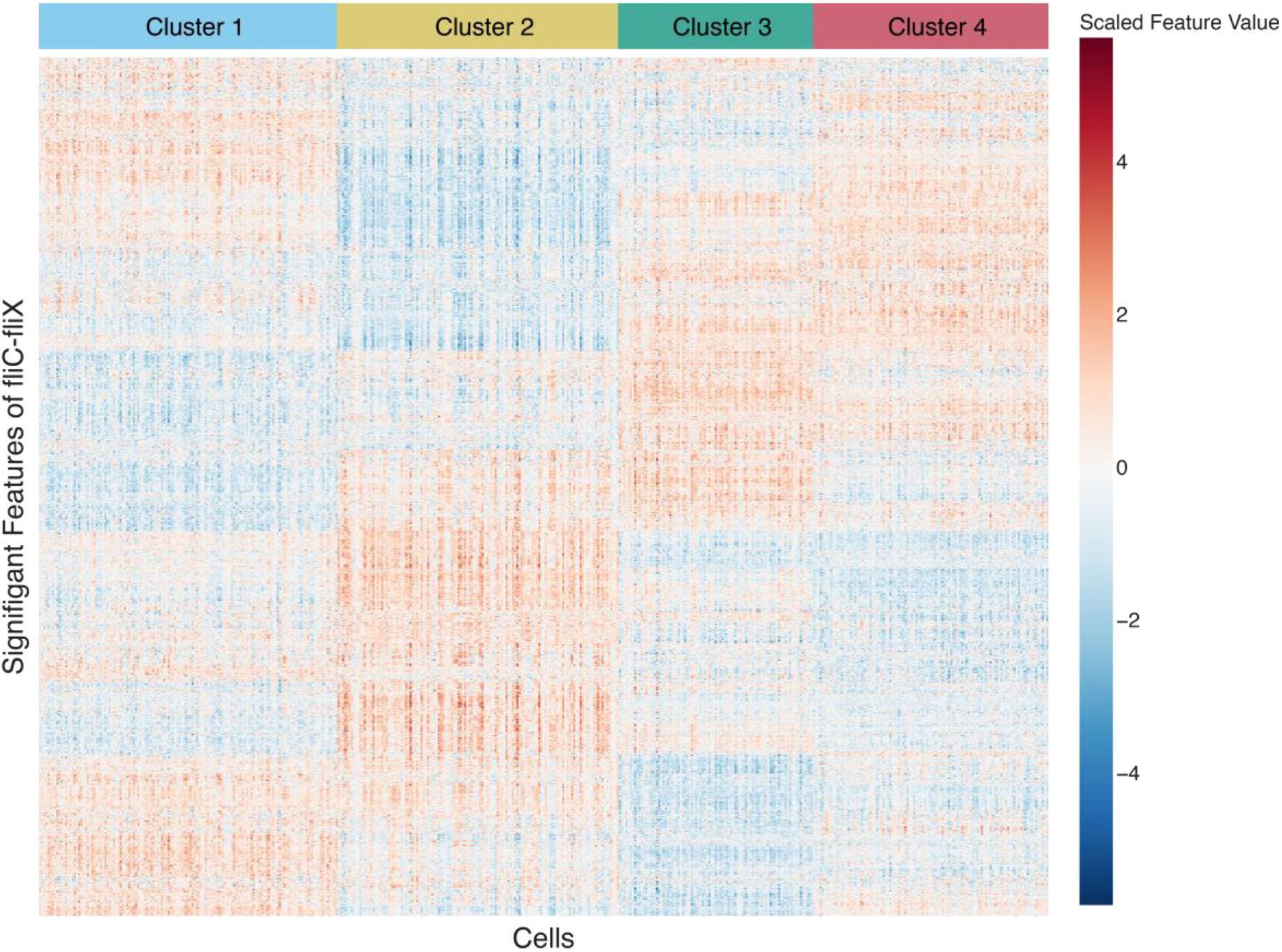
Heatmap of significant extracted UNI2-h image features for STARIT-identified clusters in standardized Bacterial-MERFISH *E. coli* data. Heatmap of scaled significant (p-value < 0.001) STARIT features for fliC-fliX, shown across the identified clusters. Rows denote extracted features, and columns denote cells.

**Supplementary Figure 14.**
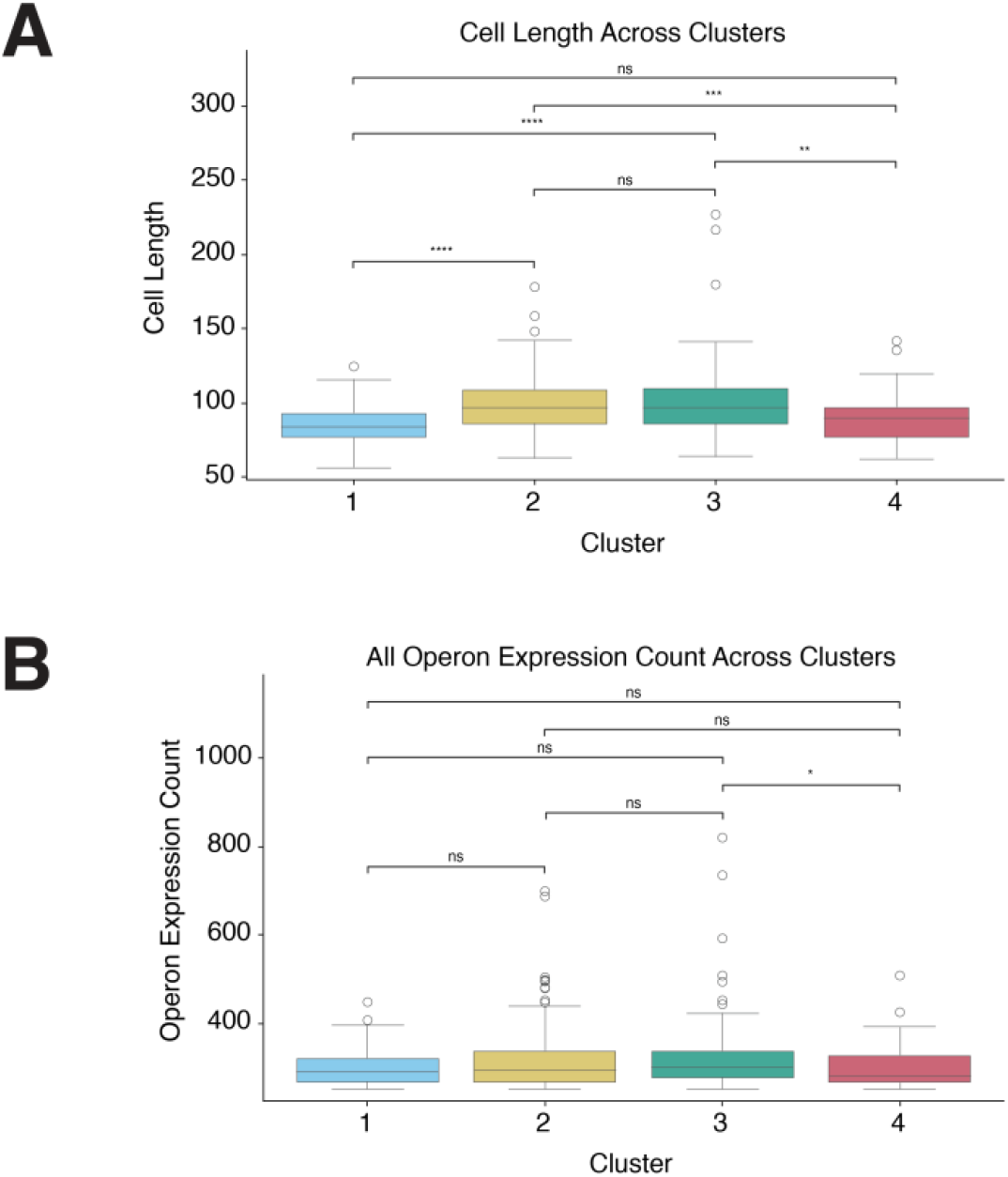
Distributions of selected expression and morphology metrics among STARIT-identified bacterial clusters. Additional boxplots showing selected supervised metrics for the four clusters identified from standardized bacterial-MERFISH *E. coli* data. (A) Distribution of total cell length in X across bacterial clusters. Boxplots display median intensity (line), interquartile range (box), and outliers, with colors denoting cluster. Statistical significance determined by Mann-Whitney U test (Bonferroni corrected). (B) Distribution of all operon expression count across bacterial clusters. Boxplots display median intensity (line), interquartile range (box), and outliers, with colors denoting cluster. Statistical significance determined by Mann-Whitney U test (Bonferroni corrected).

**Supplementary Figure 15.**
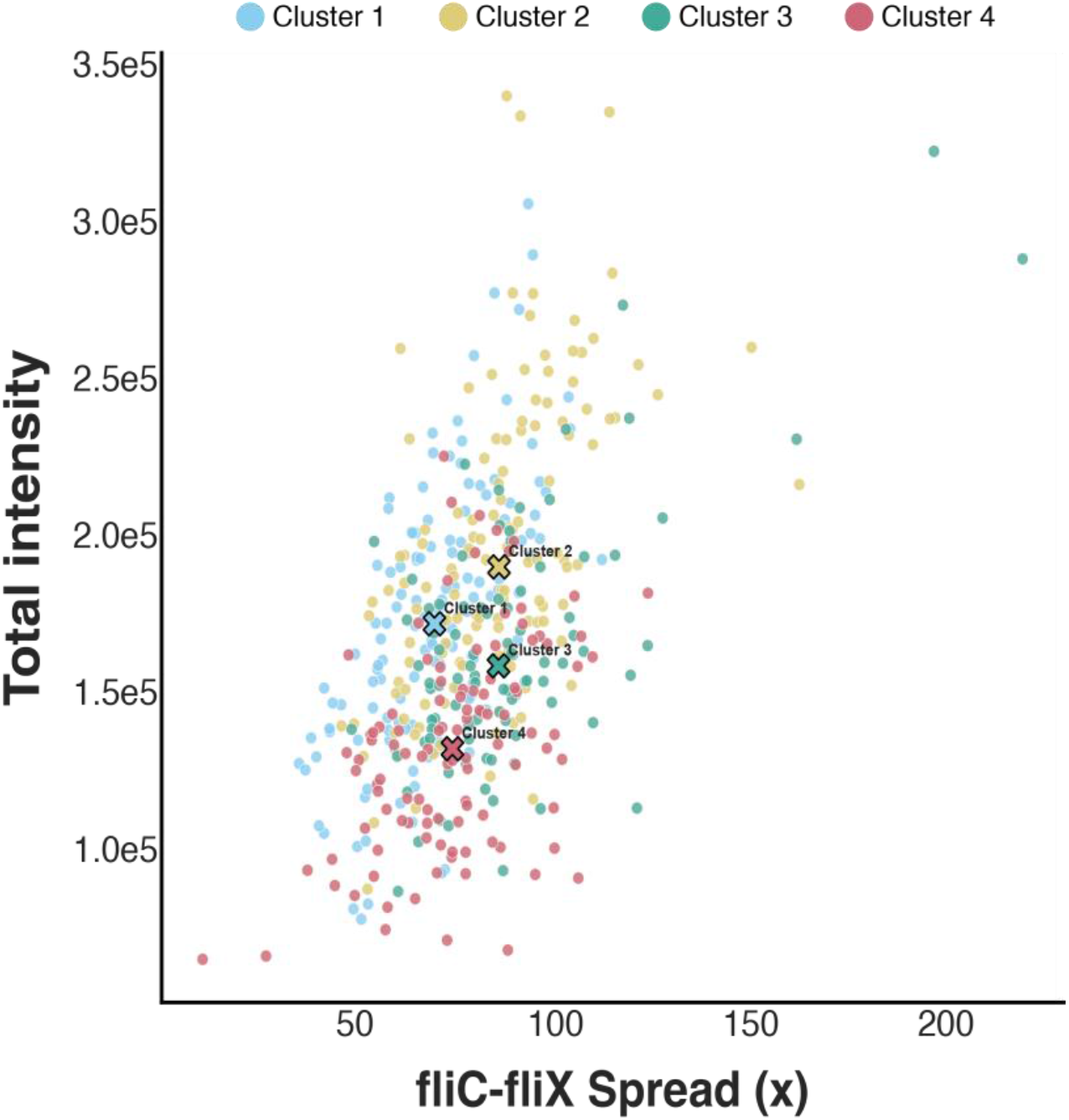
Bivariate distribution of horizontal fliC-fliX spread and total intensity among STARIT-identified bacterial clusters in standardized Bacterial-MERFISH *E. coli* data. Scatterplot of the horizontal spread of the fliC-fliX molecular distribution versus its total summed image intensity in standardized bacterial-MERFISH cells. Each point represents one *E. coli* cell and is colored by Louvain cluster assignment. Black × symbols indicate the cluster centroids in the two-dimensional metric space and are labeled by cluster.

## Notes

https://github.com/JEFworks-Lab/STARIT/tree/main/data

https://linnarssonlab.org/osmFISH/availability/

https://datadryad.org/dataset/doi:10.5061/dryad.n5tb2rc4d.

### Competing Interest Statement

The authors have declared no competing interest.

### Summary of Updates

This version has been substantially revised to include additional validation and characterization of STARIT. We added simulation studies evaluating polarized transcript localization, continuous nuclear-to-perinuclear localization trajectories, and reduced transcript capture approximating partial cell-volume sampling. We also benchmarked multiple computer-vision feature extractors, including ResNet18/50/101/152, CTransPath, and UNI2-h, demonstrating that feature-extractor choice can influence sensitivity to specific spatial properties. The bacterial-MERFISH analysis was substantially expanded by rotationally standardizing E. coli cells and applying the vision transformer pathology foundation model feature extractor UNI2-h to minimize orientation-driven variation. STARIT subsequently identified four bacterial clusters, which we further characterized using interpretable metrics, revealing differences in fliC-fliX spatial organization not explained by expression magnitude alone. We additionally expanded methodological descriptions, limitations, discussion of feature interpretability and model selection, and provided computational runtimes/specifications. The manuscript and supplementary materials have been reorganized and revised throughout for improved methodological transparency and clarity. An Author Summary and expanded data & figures have also been added.

